# Endothelial cells signaling and patterning under hypoxia: a mechanistic integrative computational model including the Notch-Dll4 pathway

**DOI:** 10.1101/2023.05.03.539270

**Authors:** Rebeca Hannah de Melo Oliveira, Brian H. Annex, Aleksander S. Popel

**Author notes:** Correspondence: Rebeca Hannah de Melo Oliveira.

## Abstract

**Introduction:** Several signaling pathways are activated during hypoxia to promote angiogenesis, leading to endothelial cell patterning, interaction, and downstream signaling. Understanding the mechanistic signaling differences between endothelial cells under normoxia and hypoxia and their response to different stimuli can guide therapies to modulate angiogenesis. We present a novel mechanistic model of interacting endothelial cells, including the main pathways involved in angiogenesis.

**Methods:** We calibrate and fit the model parameters based on well-established modeling techniques that include structural and practical parameter identifiability, uncertainty quantification, and global sensitivity.

**Results:** Our results indicate that the main pathways involved in patterning tip and stalk endothelial cells under hypoxia differ, and the time under hypoxia interferes with how different stimuli affect patterning. Additionally, our simulations indicate that Notch signaling might regulate vascular permeability and establish different Nitric Oxide release patterns for tip/stalk cells. Following simulations with various stimuli, our model suggests that factors such as time under hypoxia and oxygen availability must be considered for EC pattern control.

**Discussion:** This project provides insights into the signaling and patterning of endothelial cells under various oxygen levels and stimulation by VEGFA and is our first integrative approach toward achieving EC control as a method for improving angiogenesis. Overall, our model provides a computational framework that can be built on to test angiogenesis-related therapies by modulation of different pathways, such as the Notch pathway.

## 1 INTRODUCTION

The formation of new blood vessels from the existing vasculature, called angiogenesis, is regulated by different cells and processes (Carmeliet, 2005; Eelen et al., 2020; Zhang et al., 2022). It is generally classified as sprouting angiogenesis (sprouting of endothelial cells from a mother vessel based on environmental cues) or intussusceptive angiogenesis (vessel splitting internally into two daughter vessels), with the former being better characterized and studied than the latter, and both essential for regeneration of microvasculature (Arpino et al., 2021). Dysregulation in angiogenesis has been associated with the development and complication of different diseases, such as cancer, diabetic retinopathy, age-related macular degeneration, and cardiovascular diseases, including peripheral arterial disease (PAD) (Zhang et al., 2022). Many studies focus on modulating and controlling angiogenesis as a therapeutic approach for treating ischemic diseases and cancer (Teleanu et al., 2019; Annex and Cooke, 2021; Zhang et al., 2023). PAD, for instance, is an atherosclerotic disease characterized by lower limb ischemia; the greater the extent of the endogenous angiogenic response, the lesser the patients’ symptoms. Often accompanied by severe microvascular disease, PAD affects more than 200 million people worldwide and is associated with 53%– 90% of all major amputations of the lower limb (Londero et al., 2019; Aday and Matsushita, 2021). Currently, there is no cure for this disease, and medical therapies have limited efficacy in treating it. Therapeutic angiogenesis remains an approach that aims to alter the course of the disease (Annex and Cooke, 2021; Han et al., 2022). Among the main signaling pathways that regulate angiogenesis are the Notch signaling and the oxygen-sensing hypoxia-inducible factor (HIF) pathways. Under hypoxia, VEGFA (and possibly other angiogenic factors) is released in the hypoxic tissue, guiding endothelial cells (ECs) to the ischemic site to restore blood flow and oxygen availability. Once attracted, ECs compete for the tip position, a dynamic process guided by the Notch signaling pathway and affected by hypoxia (Figure 1) (Naito et al., 2020; Naiche et al., 2022). Additionally, EC survival under hypoxia is stimulated through autocrine VEGFA signaling, although it does not significantly affect angiogenesis (Lee et al., 2007).

**FIGURE 1.**
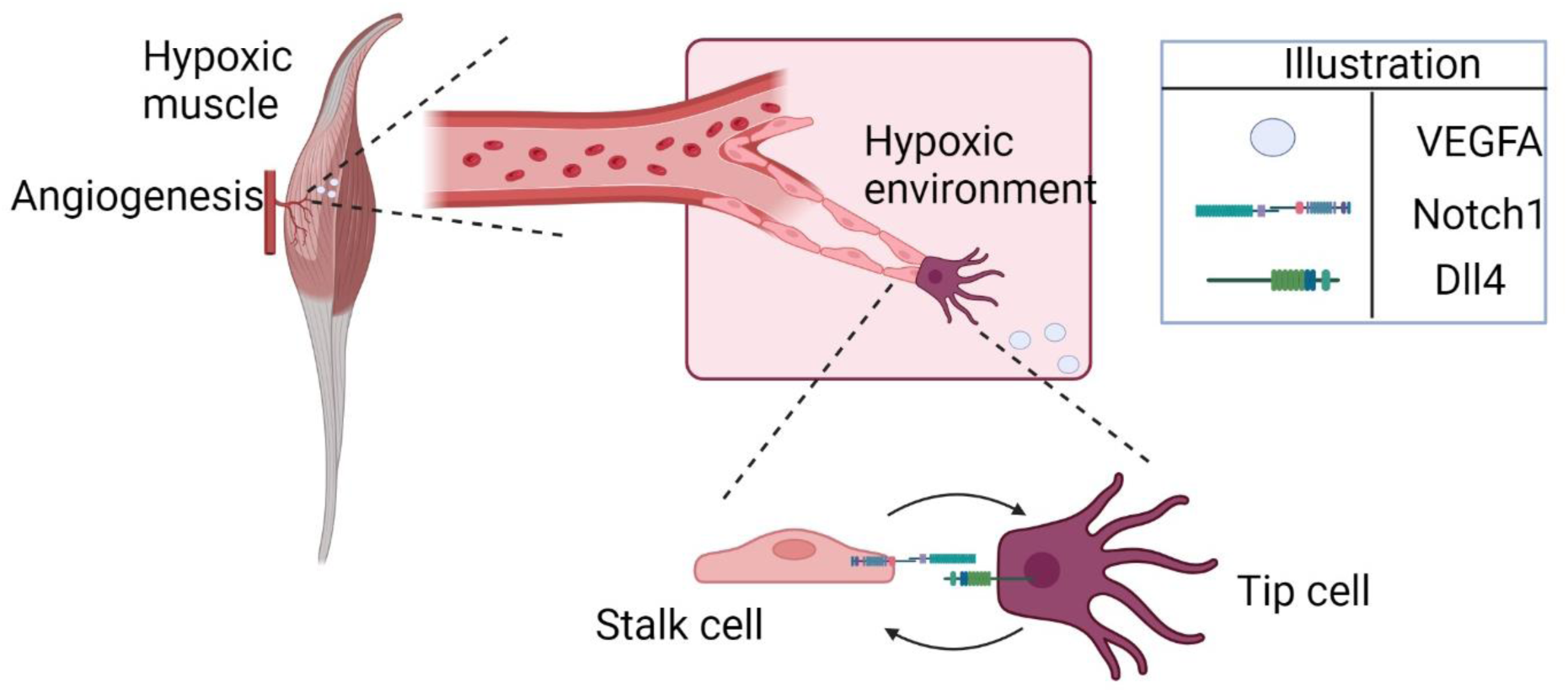
Hypoxia-induced cell patterning during angiogenesis (illustrated here for the specific example of ischemic skeletal muscle). Endothelial cells approach the hypoxic region, attracted by higher concentrations of VEGFA. The cells interact through Notch1-Dll4, competing for the tip position.

The Notch signaling pathway is a major regulator of cell fate, cell differentiation, vascular patterning, and intercellular communication in processes such as angiogenesis and tumor growth. The Notch pathway is highly conserved across vertebrates, and it affects different cells, such as macrophages (Lin et al., 2018), endothelial cells (ECs), smooth muscle cells (SMCs) (Fouillade et al., 2012), and pericytes (Tefft et al., 2022). In EC differentiation, the Notch ligands Delta-like ligand 4 (Dll4) and Jagged 1 render different effects when interacting with Notch receptors. While Dll4 is a negative regulator of tip cell formation, Jagged 1 positively regulates tip cell formation and sprouting (Benedito et al., 2009; Pedrosa et al., 2015). Dll4 and Jagged also have a role in the spatial control of sprouting (Tiemeijer et al., 2022). This competition between Notch ligands to bind to the Notch receptor is required to promote functional revascularization and tissue reperfusion.

Different cells in ischemic tissues release pro-angiogenic factors such as vascular endothelial growth factor (VEGFA) (Couffinhal et al., 1998; Rissanen et al., 2002). VEGF induces endothelial cell migration and sprouting in a chemotaxis-dependent fashion (Lee et al., 2022). This pro-angiogenesis mechanism depends on environmental cues such as low oxygen levels, driven by the HIF signaling pathway. Once low oxygen levels are sensed in the tissue, three types of hypoxia-inducible factors (HIF-1, HIF-2, and HIF-3), are activated, each composed of two subunits (α and β). The α-subunits are regulated by changes in oxygen concentration based on proteolytic degradation and transcriptional regulation. Under ischemia, HIF1-α upregulation relates to tissue inflammation, and it has been an important therapeutic target and molecule of interest in angiogenesis models (Qutub and Popel, 2006; Qutub and Popel, 2007; Qutub and Popel, 2008; Cavadas et al., 2013; Nguyen et al., 2013; Fábián et al., 2016).

The HIF and the Notch pathways are essential in the response to ischemia, and understanding the mechanistic interactions between them and other pathways that drive EC differentiation, proliferation, and stability can help develop new therapeutic strategies for diseases such as PAD. Even for converging pathways, differences in temporal regulation affect their signaling outcomes (Jin et al., 2014). Computational models have been developed and provided important insights into understanding signaling pathways and cell-cell interactions (Subramanian et al., 2022). For instance, Zhao et al. designed an ordinary differential equation (ODE)-based computational model to understand hypoxia-responsive miRNAs control of the HIF-VEGF pathway in EC (Zhao and Popel, 2015). Also, Venkatraman et al. formulated an ODE-based mathematical model to better understand how local and intracellular conditions influence EC patterning (tip or stalk EC) (Venkatraman et al., 2016). Their model showed a partial and stable tip/stalk stage with duration determined by intracellular signaling. Recently, Kuhn and Checa proposed an elegant model of the contributions of VEGF receptors to lateral inhibition during sprouting angiogenesis, integrating agent-based and ODE-based modeling (Kühn and Checa, 2019). Despite advances in experimental biology methods, observing signaling interactions between two microvascular endothelial cells under specific conditions can be challenging. Thus, *in silico* models can provide insights to help understand the mechanisms of EC patterning, signaling, and behavior under pathological conditions.

To date, most computational models of endothelial cell patterning during angiogenesis are based on Cellular Pots or agent-based modeling strategies (Kühn and Checa, 2019). Previous agent-based models on Notch signaling report how VEGF concentration relates to the elongation, migration, and proliferation of ECs (Qutub and Popel, 2009). Recently, Koon and others presented a computational model of EC patterning focusing on Notch signaling heterogeneity to explain a greater variety of cell patterning; they report hybrid endothelial cells with tip and stalk characteristics (Koon et al., 2018). A more comprehensive Boolean model, including different signaling pathways in EC during angiogenesis, was presented by Weinstein et al., incorporating the molecular regulatory network, extracellular microenvironments, and loss- and gain-of-function mutations (Weinstein et al., 2017). Mechanistic models of signaling networks in EC have also been developed. Bazzazi et al. designed a rule-based computational model implemented in BioNetGen to assess the effects of thrombospondin-1 (TSP1)-CD47 signaling through VEGF on ERK1/2 and calcium (Bazzazi et al., 2017). Using rule-based modeling, the group also investigated TSP1 inhibition of VEGF signaling to Akt-endothelial nitric oxide synthase (eNOS) (Bazzazi et al., 2018). A hybrid multi-scale model of endothelial cells during angiogenesis has also been presented by Stepanova et al., including features such as branching, cell mixing, and the brush border effect; their work indicated a dependence on the time evolution of cell mixing and the newly formed branching structure (Stepanova et al., 2021). Mechanistic computational models of intracellular and intercellular events should include the major and most relevant pathways known to the field studied (Zhang et al., 2022). Additionally, the methodology used for model building should be consistent and follow guidelines to achieve good modeling practices (Mitra and Hlavacek, 2019).

The complex molecular signaling networks that drive angiogenesis and determine EC fate are largely affected by hypoxia; note that the effect may be context-dependent and differ in different tissues such as ischemic skeletal muscle, tumor, or ischemic retina. Exogenous stimulation through VEGF and signaling between interacting ECs determine cell fate and guide downstream signaling, contributing to functional angiogenesis. Although the basic mechanisms that regulate EC patterning during angiogenesis are known, mechanistic details of the interaction between pathways and their effects and how abnormal conditions affect them need further studies (Zhou et al., 2022). In this work, we assess changes in the dynamics of such networks under hypoxia conditions to understand how VEGF, Notch, and HIF signaling pathways interact with each other and regulate downstream pathways, as well as how they affect EC patterns through time. Here we also present a comprehensive methodological approach for model development, based on recent reviews on good modeling practices, and we apply this methodology to the system of interest. Using a mechanistic model of two interacting ECs, we evaluate their signal exchange under varying oxygen conditions, investigate the effects of different interventions on cell pattern determination, and discuss their potential application in the context of angiogenesis modulation, considering the numerous effects of hypoxia on angiogenesis-involved signaling pathways (Rodriguez et al., 2021).

### 1.1 Note on nomenclature

In this work, we consider a model of two endothelial cells, with the goal of representing their interaction during angiogenesis. Initially, we stimulate one of the cells with a higher amount of VEGFA than the other to evaluate the different signals guiding their interaction. The cell that receives a higher initial stimulus is referred to in this study as the stimulated cell, first cell, or tip cell. The cell initially receiving less VEGFA is referred to as the unstimulated cell, second cell, or stalk cell. We refer generally to HIF1α and HIF2α as HIFs.

## 2 METHODS

We formulate the model network based on biological knowledge and experimental evidence and obtain the corresponding ordinary differential equations representing each biochemical reaction. Our integrative model is based on a modular structure, each module representing a biological process, such as oxygen sensing, VEGF signaling, calcium, and NO cycling, and the Notch pathway as illustrated in Figure 2. The reactions are based on mass action, Hill, and Michaelis-Menten kinetics. The model is implemented in the Matlab SimBiology software (MathWorks Inc, 2023b). The sbml and Matlab files are provided as Supplementary Material, with the initial conditions and parameter values used (Supplementary Tables ST1– ST7). A description of model components is also provided. The model was simulated using MATLAB R2023a and includes the pathways considered more relevant to our understanding of the interaction between ECs and the effect of hypoxia. The model is composed of two cells, interacting through the Notch/Dll4 pathway (to simulate lateral inhibition) and stimulated by VEGF (similar to bolus dosages). The exogenous VEGFA bolus represents exogenous gradients of VEGFA leading to EC patterning (paracrine effect of VEGFA). Under hypoxia, we also model an increase in exogenous VEGFA and in intracellular VEGFA production by the EC. Given evidence from the literature, we assume that VEGFA produced by EC under hypoxia does not affect EC patterning and angiogenesis but is present to promote cell survival signal through Akt and Nitric Oxide pathways (Lee et al., 2007). Therefore, we do not consider the transport of EC-produced VEGFA between the two interacting cells.

**FIGURE 2.**
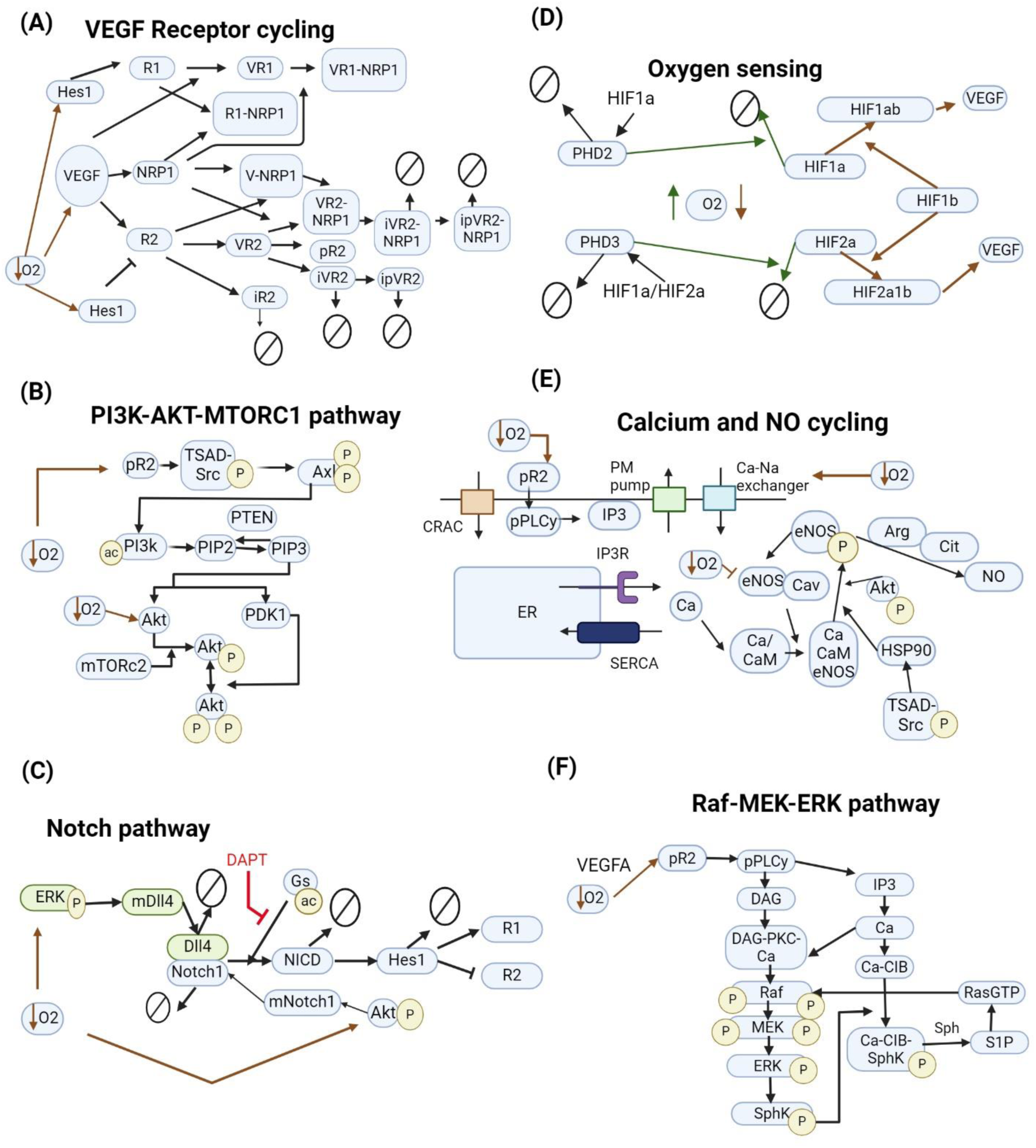
Pathways included in the model. (A) VEGF interacting with VEGFR1, VEGFR2, and NRP1. Inhibition of VEGFR2 by Hes1 and upregulation of VEGFR1. (B) Pathway leading to the downstream phosphorylation of Akt on ser473 and thr308 and activation of mTORC1. (C) Notch signaling pathway influenced by hypoxia. Green and blue colors represent different cells. DAPT, a γsecretase inhibitor, inhibits NICD cleavage by γsecretase. (D) HIF signaling pathway leads to the production of VEGFA. Solid green lines indicate normoxia-induced events. Red dashed lines indicate hypoxia-induced events. (E) Calcium cycling module includes current through the CRAC channels and PM pump, endoplasmic reticulum calcium cycling through SERCA and IP3R, and indirect stimulus of IP3 by VEGF. Na + -Ca2+ exchanger increases intracellular calcium concentration under hypoxia. (F) MAPK-ERK signaling pathway interacting with the calcium cycling module. In all diagrams, arrows pointing to other arrows represent a binding event, e.g., VEGF + R1 -> VR1.

### 2.1 Model parameterization

We use the modeling methodologies most appropriate for our model design and development (Chis et al., 2011; Villaverde et al., 2016; Jacob et al., 2023; Rey Barreiro and Villaverde, 2023). Our methodology is summarized in Figure 3. The pathways included in the model were described in the Introduction; a more detailed description is included as a Supplementary Material.

**FIGURE 3.**
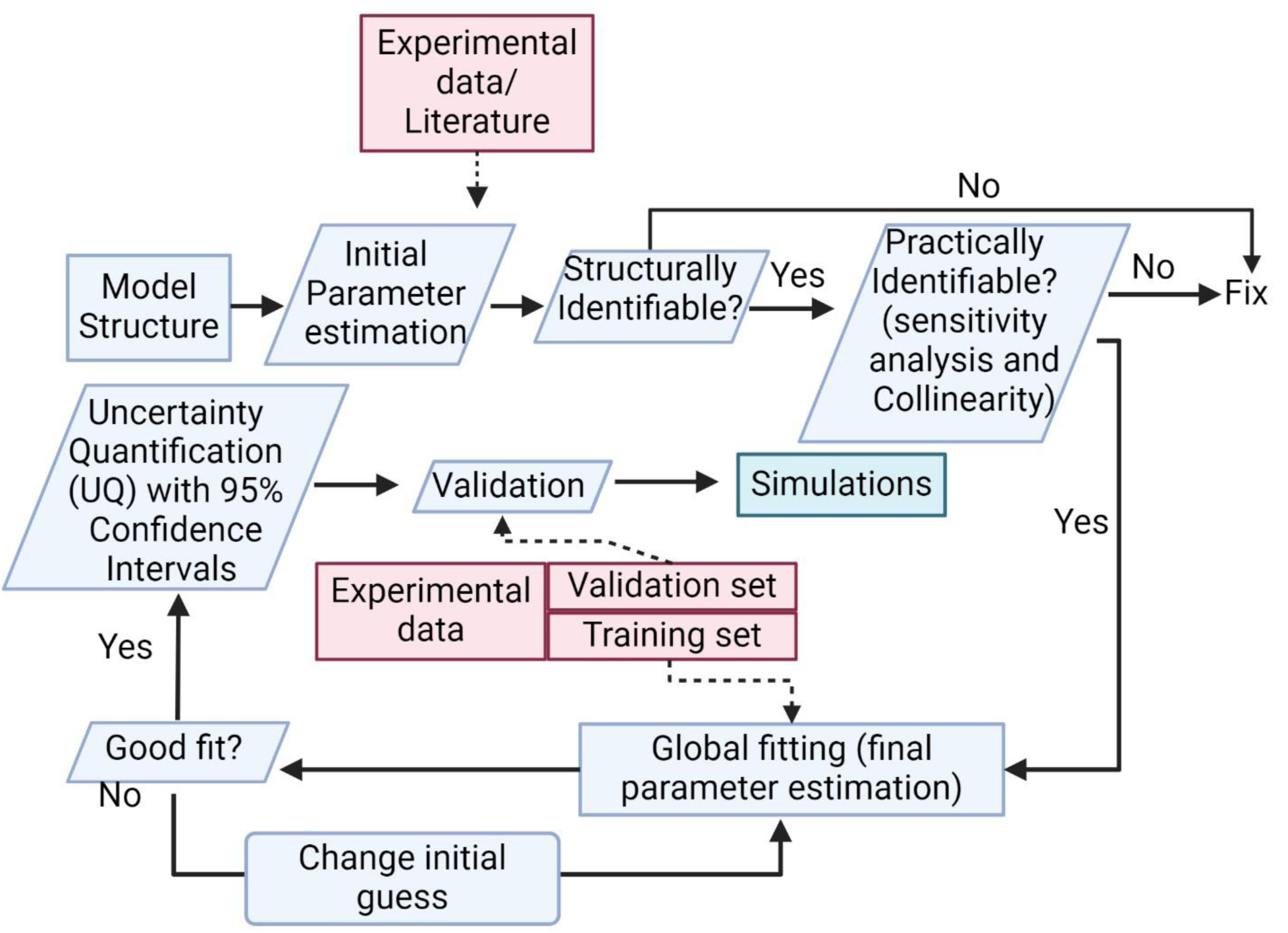
Methodology diagram. The diagram shows important steps in model development, such as structural and practical parameter identifiability, global sensitivity analysis, and uncertainty quantification.

After defining the model structure based on previous mechanistic models proposed by our group and others that follow current knowledge of the biological signaling pathways involved in angiogenesis, we proceeded with defining the initial values for species and parameters included in the model. Given our model’s high level of detail, we followed initial values presented by previous models when possible (Zhao and Popel, 2015; Venkatraman et al., 2016; Bazzazi et al., 2017; Bazzazi and Popel, 2017; Bazzazi et al., 2018; Kühn and Checa, 2019), to limit the number of parameters to be fit and reduce computational cost from optimization. These are noted in the Supplementary Table ST2, under the “Notes” column, for clarity. We work with units of concentration (μM), so values may have been converted between models. The units of time-dependent parameters are given in either seconds or hours. More information on unit conversion is presented in the Supplementary Material section.

From our initial estimates and model structure, we follow a pre-defined methodological approach for model development based on the following steps: I) Define model structure → II) Structural and practical identifiability analysis → III) Global optimization (fitting of parameter values to approximate experimental data) → IV) Uncertainty Quantification (UQ) and Validation. Next, we detail the methods used for each step.

#### 2.1.1 Define model structure

Our model structure integrates some of the major pathways included in sprouting angiogenesis under hypoxia in EC, i.e., VEGF/VEGFR, Dll4/Notch, HIF, Calcium, and the phosphorylation of eNOS, ERK and Akt (Zhang et al., 2022). A detailed description of the modeled pathways is provided in the Supplementary Material.

#### 2.1.2 Structural identifiability analysis (SIA)

SIA investigates the possibility of obtaining a unique value for each parameter in a model considering a known and theoretically perfect model structure. It allows us to select which parameters can be considered for fitting. Several methods are available for performing this analysis, as recently reviewed (Chis et al., 2011; Rey Barreiro and Villaverde, 2023). Several methods exist to perform SIA, and at least two are currently compatible with Matlab interface, GenSSI and STRIKE-GOLDD. STRIKE-GOLDD 4.0 provides the option of model automatic reparameterization (Massonis et al., 2023) and an optimized algorithm for computationally expensive rational models (Díaz-Seoane et al., 2023). GenSSI 2.0 provides written reports of structural identifiability analysis as well as a graphic representation. It integrates identifiability tableaux with generating series approach to identify the uniqueness of solutions to an estimation problem (Chis et al., 2011; Ligon et al., 2018). As the two methods have their advantages, we initially performed SIA with both. However, we found that GenSSI 2.0 performed better in terms of time. Therefore, our SIA results are presented graphically, simulated with GenSSI. The SIA results indicate which parameters can be used as input in the model fitting (identifiable) and which have to be obtained from literature sources or assumed (non-identifiable). Our SIA results are presented in Supplementary Figure S1. We include global sensitivity analysis using PRCC to evaluate how the assumed parameters affect the model outputs (Supplementary Figure S4 in the Supplementary Material).

#### 2.1.3 Practical identifiability analysis (PIA)

To test if the fitted parameters can be uniquely determined from the data available, we perform PIA, to obtain confidence intervals for the parameter values. Practical non-identifiability can be caused by the absence of influence of the parameter investigated on the model observables (outcomes) or by the parameters being interdependent. We implement a method previously described to investigate PIA, through sensitivity analysis and collinearity of the sensitivities of parameters (Gábor et al., 2017). Our approach for PIA is to first perform global sensitivity analysis using Partial Rank Correlation Coefficient (PRCC), as described previously (Renardy et al., 2021), investigating which of the structurally identifiable unknown parameters are most relevant to changes in the observables of our model. Parameters with no or too little effect are considered non-identifiable. We set the cut-off values as having a PRCC greater than 0.2 or/and a *p*-value less than 0.05. Following this, we perform collinearity analysis. Our method for collinearity analysis is a straightforward elimination of parameters with opposite effects on reactions (e.g., forward and reverse reaction rates; the phosphorylation and dephosphorylation rates). For the set of unknown and non-identifiable parameters after PIA, we assume values based on the literature or previous models. The unknown and identifiable parameters are fitted based on the strategy explored in step III.

#### 2.1.4 Global optimization (GO)

To find the unknown parameter values based on experimental data available, we collected data from the literature showing species responses to certain stimuli. Specifically, we searched for time course data showing the effect of 1% O_2_ (hypoxia, equivalent to approximately 8 mmHg of O_2_ partial pressure) on species included in our model, as well as responses of different pathways to VEGFA165a stimulation. In our model, we do not consider other isoforms of VEGFA, or other growth factors, for simplification. Most data available and used in our GO analysis is from Western Blot (WB) studies, a semi-quantitative approach. As absolute values are not measured in WB, we normalize all data to its maximal concentration prior to fitting. Similarly, we use our simulations’ normalized responses (to the maximal concentration during a certain time) to fit our model parameters to the time-course dynamics of gene and protein expression. This is a standard procedure that has been used in previous models (Bazzazi et al., 2017; Bazzazi and Popel, 2017; Bazzazi et al., 2018). With the data collected and normalized, we then implement global fitting on the unknown identifiable parameters selected from SIA.

We use particle swarm optimization, available in SimBiology Model Analyzer, as used by others to evaluate mechanistic systems biology models (Song and Finley, 2020; Song et al., 2023). PSO works by using multiple candidate solutions (particles forming a parameter set) for the optimization problem. At each iteration, the algorithm investigates the parameter space, and each candidate solution registers and keeps track of its personal optimum solution and of the best solution of the entire population of solutions, using an objective function such as the weighted sum of squared residuals. The algorithm aims to minimize this function to identify the best set of optimal parameter values. We implement PSO setting the bounds of the estimated parameters as one order of magnitude above and below the baseline values assumed initially based on the literature.

On our first trials of working with PSO, the computational cost and time spent on fitting presented themselves as impractical, leading us to a different fitting strategy. In this strategy, as the data collected for model calibration can be separated into two groups (stimulation by VEGF or stimulation by hypoxia), we use this to divide the parameters to be fit into two groups as well (according to the sensitivity analysis of the observables performed previously). We classify each parameter as involved in one of 2 groups. We then start by fitting the parameters classified as more relevant to the VEGF-stimulated group and obtain their optimized values. We assign the new estimated values to the model and fit the next set of parameters (for hypoxia-related classification). Then, we test the final fitted model by comparing the responses to experimental data and through UQ. This strategy significantly reduced the time required for fitting and showed good calibration. We apply the Runs test on calculated residuals to assess the goodness of fit, as described in previous studies (Motulsky and Ransnas, 1987; Bujang and Sapri, 2018). The Runs test results are included in Table 1. To perform the Runstest, we use the Matlab function h = *runstest (x)*, where h represents a test decision for the null hypothesis that the values in *x* are in random order. For h = 1, the test rejects the null hypothesis, and zero otherwise. The test also returns a *p*-value, which represents the probability of observing a test statistic as extreme as (or more than) the value observed under the null hypothesis. In other words, a low *p*-value disputes the validity of the null-hypothesis (MathWorks Inc, 2023a).

**Table 1:** Runs Test on residuals post-fitting.

| Condition | Observable | h | p |
| --- | --- | --- | --- |
| Normoxia | peNOS | 0 | 1 |
|  | NO | 0 | 0.4 |
|  | Ca | 0 | 0.085 |
|  | VEGFR2 | 0 | 1 |
|  | pVEGFR2 | 0 | 0.8 |
|  | pAkt | 0 | 0.809 |
|  | pERK | 0 | 1 |
|  | mDII4 | 0 | 0.6 |
|  | NICD/Notch1 | 0 | 0.666 |
|  | NICD | 0 | 0.8 |
|  | Hes1 | 0 | 0.666 |
|  | pPLCy | 0 | 1 |
| Hypoxia | mVEGFA | 0 | 0.666 |
|  | HIF1a | 0 | 0.05 |
|  | HIF2a | 0 | 0.09 |

#### 2.1.5 Uncertainty quantification (UQ) and validation

UQ allows us to estimate a degree of uncertainty in our model predictions (Mitra and Hlavacek, 2019). Different methods can be applied to perform UQ, as recently reviewed and compared (Rey Barreiro and Villaverde, 2023). In this work, we assess UQ through the confidence intervals of predictions based on the fitted parameters. We implement UQ based on a Bootstrapping approach (with 100 samples, to limit computational cost, 95% confidence intervals), available in the SimBiology Model Analyzer toolbox. We present our results in Supplementary Figures S2, S3 of the Supplementary Material. We define the prediction as valid if the majority of its points fall within the 95% confidence interval. We performed several rounds of fitting altering the initial guess (using values within 2 orders of magnitude above or below the initial guess). Our model predictions use the best fit we found among those (Figures 4, 5). For model validation, we use a different dataset than the one used for model training (fitting) (Bruns et al., 2010; Park et al., 2010; Chen et al., 2013; Ubezio et al., 2016). We compare the simulated response to the experimental data points reported, as well as the confidence intervals reported experimentally (Figure 6).

**FIGURE 4.**
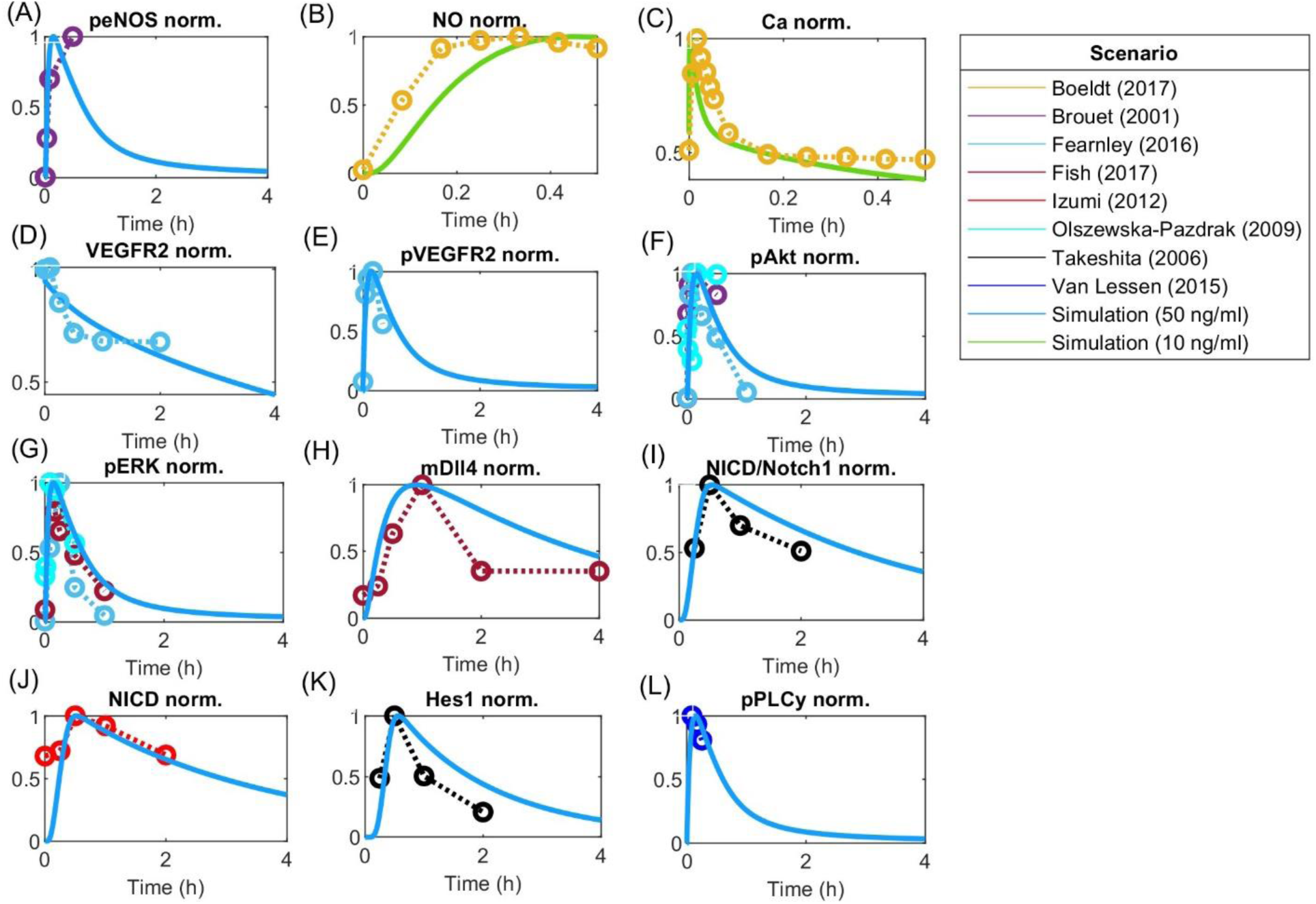
Model responses to VEGF stimulation fitted to experimental data. Global optimization with particle swarm optimization is used to fit unknown identifiable parameters to various data sets as indicated by the scenario box. Fitting was matched to VEGFA initial stimulations employed in the experimental data. Concentration is normalized to maximum value before fitting responses for (A) Phosphorylated eNOS, (B) Nitric Oxide (NO), (C) Calcium++ (Ca), (D) Surface VEGFR2, (E) Phosphorylated VEGFR2, (F) Phosphorylated Akt, (G) Phosphorylated ERK1/2, (H) Dll4 mRNA, (I) NICD relative to Notch1, (J) NICD, (K) Hes1, and (L) phosphorylated PLCy.

**FIGURE 5.**
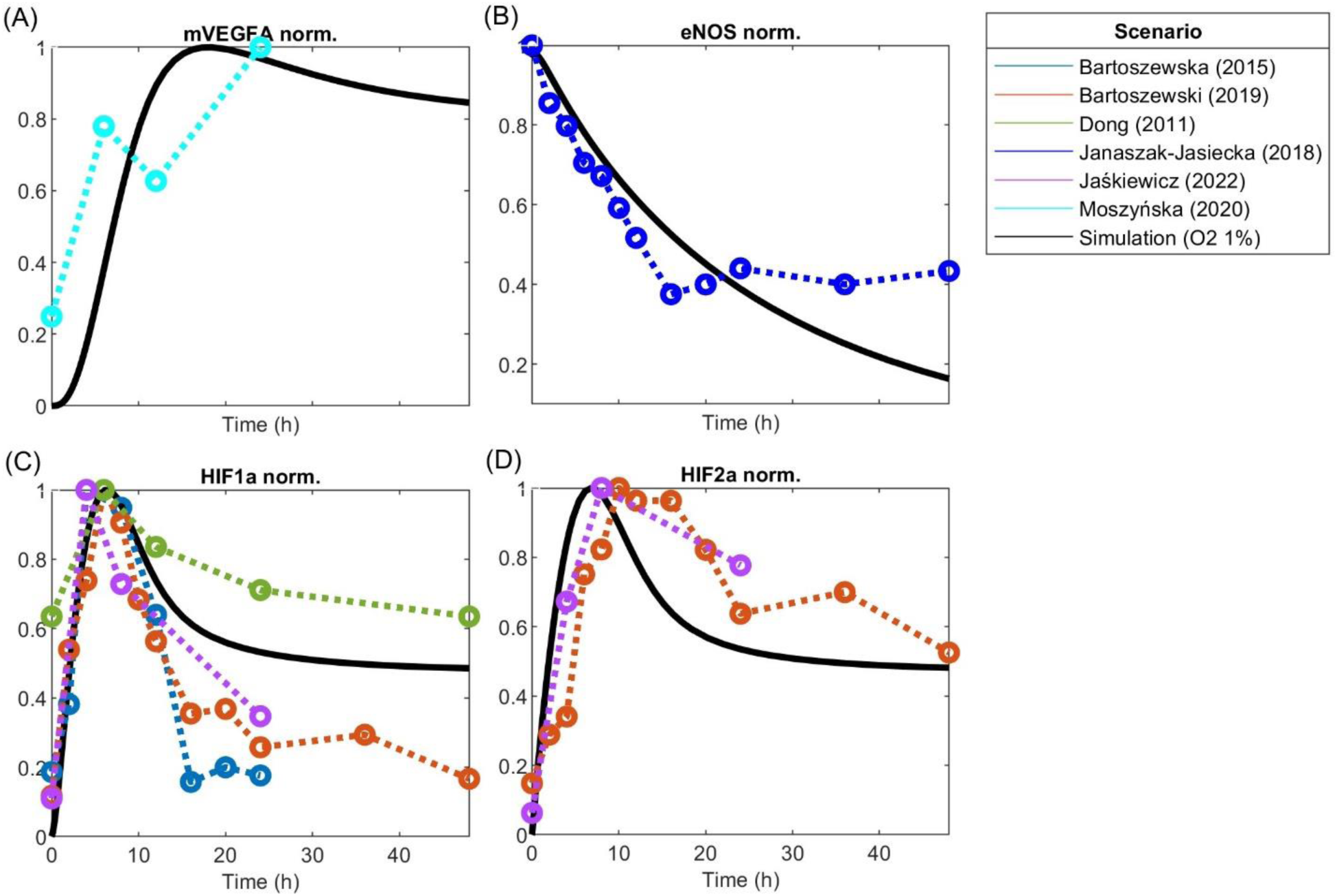
Model responses to hypoxia stimulation fitted to experimental data of hypoxia post-optimization using Particle Swarm Optimization of (A) VEGFA mRNA, (B) eNOS, (C) HIF1a protein, and (D) HIF2a protein. Concentration is normalized to its maximum before fitting.

**FIGURE 6.**
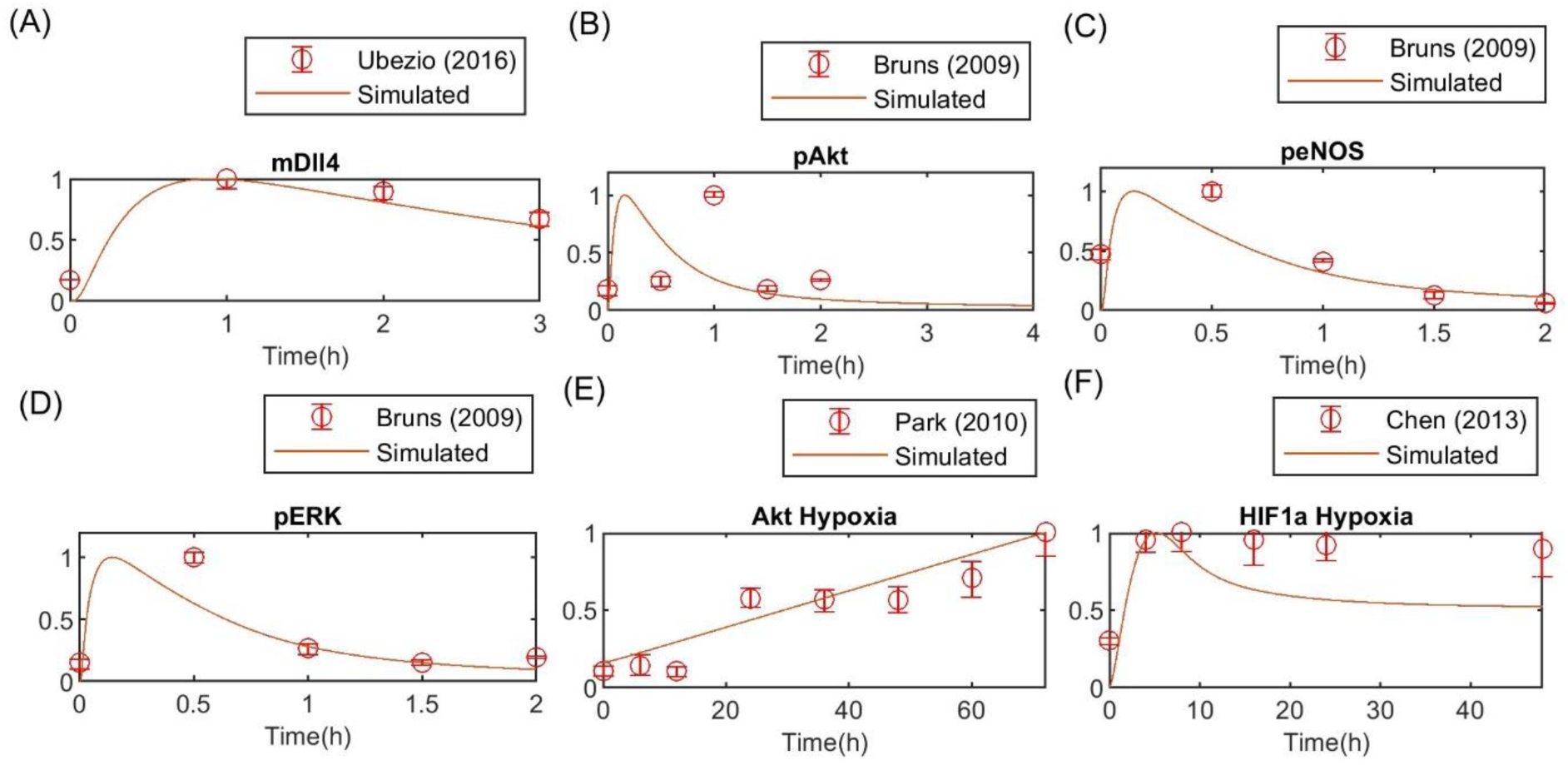
Model validation against independent experimental data for (A) Dll4 mRNA, (B) phosphorylated Akt, (C) phosphorylated eNOS and (D) phosphorylated ERK1/2, (E) Akt response to hypoxia, (F) HIF1a response to hypoxia. Concentration is normalized to the maximum value to match experimental data.

After performing this methodology for model design, calibration, and validation, we proceed to our simulations to investigate cell-cell interaction and the effects of hypoxia and Notch signaling.

### 2.2 Endothelial cell pattern index and global sensitivity analysis

As one of our goals with this model is to investigate the patterning behavior of two endothelial cells interacting under hypoxia, we define the Endothelial Cell Pattern Index (ECPI) as follows:

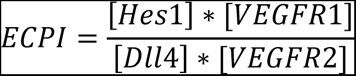

The index represents the ratio between markers of stalk cells and tip cells. In general, Tip cells are known to express higher levels of Dll4 and VEGFR2, while a higher level of Hes1 and VEGFR1 characterizes stalk cells (del Toro et al., 2010; Xu and Li, 2022). Therefore, the ECPI is a dimensionless index representing the ratio of molecular expression seen in Stalk cells relative to what is seen in Tip cells during sprouting angiogenesis. To evaluate the effect of different model parameters and species on the index, we perform a global sensitivity analysis of ECPI, described next.

Global Sensitivity Analysis (GSA): with the finalized model structure and parametrization, we proceed with the GSA of the ECPI. We implement PRCC based on the algorithm from Renardy et al. (2019), evaluating the sensitivity of PRCC in the two simulated interacting cells at two pre-specified time points (1h and 12 h) under hypoxia (1% O_2_) and normoxia (21% O_2_) (Renardy et al., 2019). The choice of time points is based on the expected higher concentration of molecules included in the ECPI given VEGF stimulation. In all cases, the parameter values are varied by 1.5 (lower bound = initial value/1.5; upper bound = initial value * 1.5).

Additionally, to investigate the influence of VEGFR2 and VEGFR1 initial concentrations on cell patterning, we performed local sensitivity analysis of VEGFR2 and VEGFR1 on Hes1 in both cells. The results are reported in Supplementary Figure S5.

### 2.3 VEGF gradient between neighboring endothelial cells

EC patterning depends on VEGFA gradient between cells, with a higher VEGFA directing the cell to assume a tip position in the sprout, while the neighbor cell assumes a stalk position due to lateral inhibition by the Notch pathway (Gerhardt et al., 2003). To perform our simulations with cell patterning, we calculate the gradient of VEGF between cells based on their length when fully extended and the length of filopodia. We consider that there is a decrease in VEGF of 3.5% for every 10 μm (Ji et al., 2007), and the cells are about 100 μm of length (Adamson, 1993) and filopodia has about 10 μm of length (Ucla et al., 2022). We also assume that most VEGF is being sensed at the end of the filopodia and VEGF concentration has an initial value of 0.0012 μM. Additionally, we assume that VEGF receptors are distributed along the length of the stalk cell, so we estimate VEGF captured by the stalk cell as an average based on its length. Figure 7 shows the diagram of our assumptions. Based on that, we calculate the VEGF sensed by each cell as 0.0012 μM for the first cell, and 0.0006 μM for the second cell (average between 0.00075 μM and 0.000487 μM).

**FIGURE 7.**
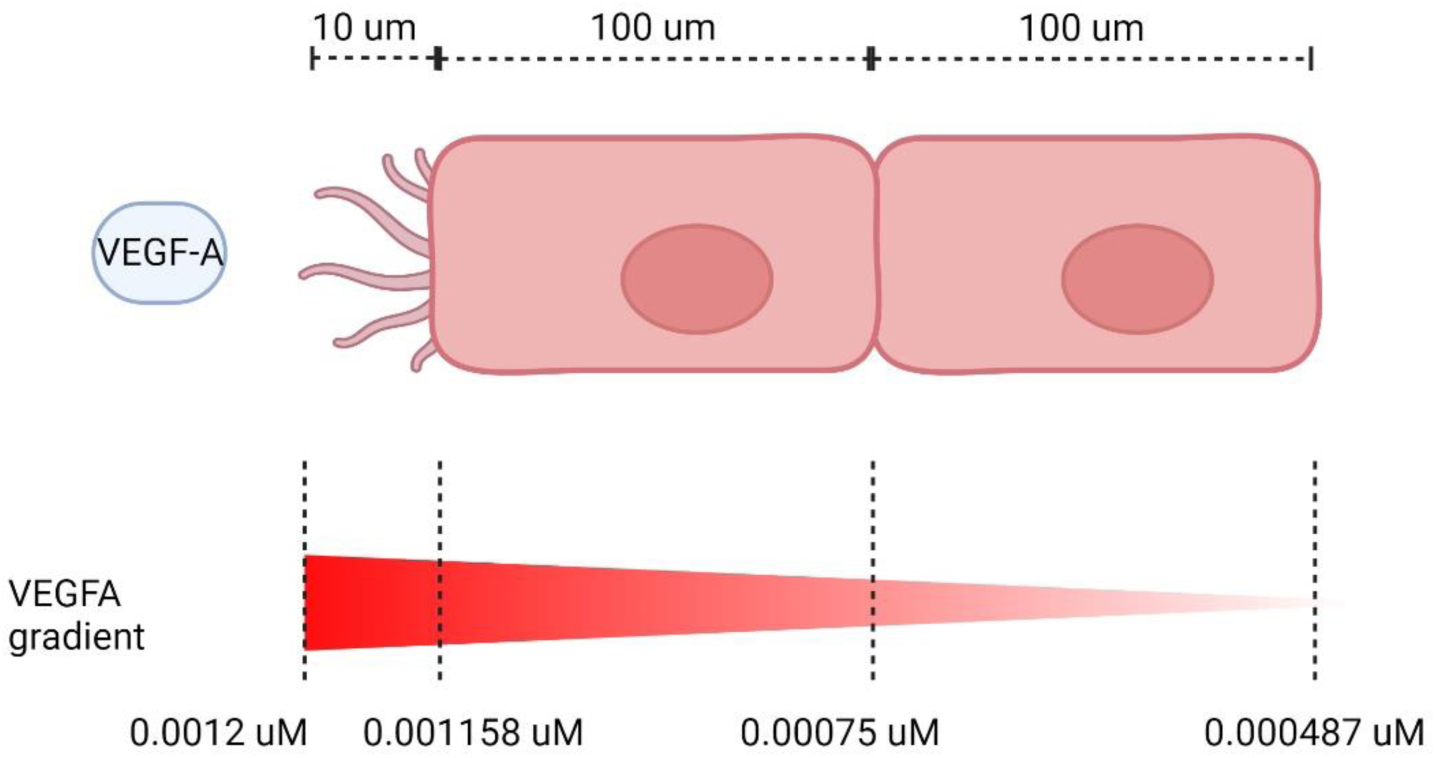
VEGF gradient on neighboring ECs. Dimensions not to scale.

## 3 RESULTS

In this section, we present the results of our model construction, methodological approaches, and simulations.

### 3.1 A structured methodology for large mechanistic model development

Mechanistic computational models describing intracellular and intercellular cell signaling under different conditions require an increasing number of reactions, parameters, and state variables to represent biological conditions more realistically. As the number of components increases, so does the complexity and computational cost for fitting and performing simulations. Good modeling practices are also required to obtain trustable models with reasonable accuracy. With that in mind, in this work, we present a structured methodology for the development of a large mechanistic model of EC signaling during angiogenesis, following the steps described in the *Methods* section (Figure 3).

As computational cost and simulation time are two factors that modelers wish to optimize, we propose that the first step following the definition of the model structure is identifiability analysis. By using structural and practical identifiability analysis we can define the parameters that can theoretically have unique values given the data used and model structure, while also reducing the number of parameters to fit. For this step, we tested different methods suggested in the literature and found the ones that worked best in terms of compatibility with Matlab and model size. The next step to limit computational cost that we use is to perform model fitting with Pattern Search, dividing the observable data set and parameters by groups (in our case, two groups: normoxia and hypoxia). To find one final set of parameters to perform the simulations, we perform fitting to find parameters for one of the groups, and then we set the values found as base values, prior to fitting to the data of the second group. By doing so, we significantly reduce the computation time for fitting (15-fold). Finally, we perform global sensitivity analysis on all parameters in the model, to find those that are more influential to a specific observation (i.e., the ECPI of each cell).

### 3.2 Model structure and parameterization

The integrative model was designed based on established knowledge regarding the included models and on previous mechanistic computational models (Zhao and Popel, 2015; Venkatraman et al., 2016; Bazzazi et al., 2017; Bazzazi and Popel, 2017; Zhao et al., 2017; Bazzazi et al., 2018; Bazzazi et al., 2018; Wu and Finley, 2020; Jaśkiewicz et al., 2022; Ferrante et al., 2023) of EC signaling, patterning and hypoxia. Figure 2 shows the pathways included in the model.

Overall, the model describes the regulation and coordination by Notch and HIFs of the cell interactions and downstream signaling. The model comprises two representative cells, initially configured similarly and later stimulated by different concentrations of VEGF and is divided into 6 modules: the Notch pathway, the oxygen sensing module, the VEGF-VEGFR pathway, the Akt-eNOS pathway, the Raf-MEK-ERK, and the calcium and NO cycling module. The final model comprises 183 species (including Vext), 222 reactions, 184 parameters, and 183 ODEs (including Vext and considering HIF1β constant in both cells and Jcrac as a state variable in both cells). Initially, we had 66 unknown parameters. On these, we performed identifiability analysis (SIA and PIA) to determine which were identifiable and which required assumptions or estimations from the literature. Our results from SIA and PIA are included as Supplementary Material [Supplementary Figure S1 (SIA, equivalent parameter names listed in Supplementary Table ST2), Supplementary Figure S6–13 (PIA, sensitivity of observables)]. As previously discussed, on PIA, we also exclude parameters that are collinear (i.e., parameters with opposite effects in the same reaction, keeping only one of them). Finally, we found 20 parameters that could be fitted. The values of the remaining non-identifiable parameters were either estimated from ranges in the literature or assumed. These are indicated as such in the Supplementary Material (Supplementary Table ST2). Since many unknown parameters were part of the Notch signaling pathway, considered crucial in this work, we performed PRCC to investigate their effects on the ECPI. The results are shown in Supplementary Figure S4. As expected, some parameters in the Notch pathway have a high influence on the ECPI, and, therefore, influence cell patterning; specifically, the degradation rates of NICD, Hes1, and Dll4 mRNA, the multiplication factor for the expression of HES1 (dependent on NICD cleavage) and Dll4 mRNA (dependent on ERK1/2 phosphorylation), and the translation rate of Dll4. These are kept fixed for the simulations, and the results presented here regarding patterning depend strongly on their values.

To determine model identifiable parameters, we fit and validate the model to experimental data from human endothelial cells (HUVECs) and, if not available, other EC types, available in the literature (Figures 4–6). All data simulated were normalized to their maximum concentration for comparison with literature data. We obtained the initial absolute concentrations in units of µM from previous mechanistic models of endothelial cells and data from the literature. Initial VEGF receptor values were estimated as 6000 VEGFR2/cell and 2000 VEGFR1/cell (Imoukhuede and Popel, 2011), considering a cell area of 1000 µm^2^, with 1 * 10^−12^ L volume (Bazzazi et al., 2017). Neuropilin-1 density per cell has been estimated in the range of 10^3^−10^6^ number per cell (Mac Gabhann and Popel, 2006).

Starting at an initial estimate of 25,000 receptors per cell (Soker et al., 1996), we hand-tune the initial concentration of [Neuropilin1], for both cells, obtaining 0.0664 µM as the initial molar concentration, equivalent to about 40,000 receptors per cell in both cells, a value smaller than estimated in other works (Imoukhuede and Popel, 2011). Quantifying an absolute number of surface receptors in specific cell types can be challenging, despite recent efforts (Chen and Imoukhuede, 2019; Sarabipour et al., 2024), and should be adapted in models as needed. We evaluate this aspect by inspecting the sensitivity of Hes1 in both cells to changes in VEGFR1/2 initial concentration, as an additional analysis (Supplementary Figure S5). To evaluate the effect under patterning, we stimulate the cells with different amounts of VEGFA by activating Vext (for normoxia and hypoxia). Our results indicate that time and O_2_ level influence how much the initial amount of receptors influences Hes1. As expected, VEGFR2 has a higher influence than VEGFR1 in all simulated scenarios, although we can still see some influence of VEGFR1 during initial time points (2 h). Under normoxia and hypoxia, the model indicates a similar effect for the 2-h simulation, but for the 48-h simulation VEGFR2 of the second cell exerts a higher influence on Hes1 of the first cell under normoxia than hypoxia, and VEGFR2 of the first cell poses a stronger effect on Hes1 of the second cell under hypoxia.

To show cell differentiation based on higher VEGF sensing under normoxia, all species concentrations are considered initially the same for the two interacting cells. To simulate cell patterning, we consider an initial stimulus of 0.0012 µM of [VEGF] for the first cell, while the second cell is set to a 0.0006 µM stimulus. This different stimulation allows us to analyze the Notch signaling pathway effects on EC patterning. As absolute concentrations for Notch1 receptors on the surface of EC during angiogenesis are not available, we estimated the initial concentration to be about the same as that of [VEGFR2] receptors (0.0099 µM). [mRNA Dll4], [mRNA Notch1], and protein [Dll4] initial values were zero for both cells.

The model is fitted under normoxia (21% O_2_), with VEGF stimulation (50 ng/mL/10 ng/mL), and hypoxia (1% O_2_, equivalent to approximately 8 mmHg of O_2_ partial pressure). The data used for calibration and validation amount for more than 150 data points from 18 different studies presenting time-course data. Given the amount of stimulation by VEGF given to each cell, we perform and present the fitting to the species in the first cell, stimulated with VEGFA 50 ng/mL (time course data for all species except NO and Ca++) or 10 ng/mL (time course data for NO and Ca++). We also present the calculated 95% confidence intervals for the predictions (as described in the Methods section) in Supplementary Figures S2, S3. We present the Runs test of the residuals calculated as a representative of the goodness of fit in Table 1. In all cases the Runs test indicates that the null hypothesis of randomness is not excluded (h = 0), and the *p* values closer to 1 indicate that there is less doubt in the validity of the null hypothesis. Our model closely follows the expected time courses seen experimentally and reproduces the expected time points at about 74% and 40% of the time (points within confidence interval over the total number of points) for the normoxia and hypoxia predictions, respectively. Although not all points fall within the prediction confidence interval, the trajectories reproduce well the expected behaviors. Additionally, using the Gaussian method to calculate the 95% confidence intervals, we find wider intervals, with more points being within them (∼90% of the time, data not shown). This difference can be due to sample size and variability of data used. As our predictions are within reasonable distance from the experimental points, we employ the fits obtained on our simulations.

#### 3.2.1 VEGF-driven pathway under normoxia

Using global Particle Swarm Optimization, we initially fit the model under normoxia conditions ([O_2_] = 209 µM) for stimulation with 50 ng/mL of VEGFA. Figures 4A–G, 5A–D present the time dynamics of the fitted species compared to the experimental data. For model validation, we show our simulations compared to data reported in two different datasets than those used for model training (Bruns et al., 2010; Ubezio et al., 2016). The results are presented in Figures 6A–D. Given few time-course data available for fitting and validation of the model (especially time-course data for state variables in the Notch pathway), we consider such initial results sufficient to evaluate the results considered in this study. As more data becomes available, the model fitting can be re-assessed following the methodology proposed in this work.

The time dynamics of Notch-related pathways (compared to data used for fitting) are shown in Figures 4H–K. Endothelial cells stimulated by VEGFA present an increase in the activation of the Notch signaling pathway, increasing NICD and Hes1 expression, which peak during the first hour of stimulation, and then decay (Takeshita et al., 2007; Izumi et al., 2012). Additionally, VEGFA stimulation is known to increase the expression of Dll4 mRNA through the ERK pathway (Fish et al., 2017).

Other events stimulated by VEGFA include the phosphorylation of VEGFR2 (panel E), Akt (panel F), and ERK1/2 (panels G) (Olszewska-Pazdrak et al., 2009; van Lessen et al., 2015; Fearnley et al., 2016). Given VEGFR2 phosphorylation, the calcium signaling pathway is activated and influences eNOS phosphorylation (panel A), Nitric Oxide production (panel B), and Calcium intracellular upregulation (panel C) (Brouet et al., 2001; van Lessen et al., 2015; Boeldt et al., 2017). For Ca++, we see a peak followed by a low persistent signaling, similar to the reported previously for cells stimulated with higher [*VEGF*](>5 ng/mL) (Noren et al., 2016). The rapid phosphorylation of VEGFR2 causes the decrease of the free surface receptors, as shown in panel D (Fearnley et al., 2016).

#### 3.2.2 Hypoxia-driven pathway

Under hypoxia ([O_2_] = 9.9 µM), several events occur in endothelial cells, as previously discussed. Given our interest in better understanding EC patterning under hypoxia, we calibrate the model parameters to data from endothelial cells under hypoxia. The results are presented in Figure 5. Once again, we compare the fitted model response to additional datasets (Figures 6E, F). The species behavior over time under hypoxia fits well the experimental data reported by others (Park et al., 2010; Chen et al., 2013).

Endothelial cells under hypoxia sense the low oxygen levels through the HIF1/2α signaling pathway, which will lead to an increase in the concentration of HIF1α (panel C) and HIF2α (Panel D) (Dong et al., 2011; Bartoszewska et al., 2015; Bartoszewski et al., 2019; Jaśkiewicz et al., 2022) followed by downstream release of VEGFA mRNA (panel A) (Moszyńska et al., 2020). Hypoxia also leads to a decrease in free eNOS levels (panel B). Limited data are available for the time course of proteins from the Notch signaling pathway under hypoxia. However, experiments by Patel and others showed an increase in Dll4 under hypoxia (Patel et al., 2005).

#### 3.2.3 Cell ECPI depends on VEGF stimulation, oxygen levels, and time under hypoxia

To observe the major pathways and parameters affecting cell patterning, we perform a global sensitivity analysis of all model parameters on the ECPI of the first and second cells. Due to a large number of parameters, we present the PRCC results for each pathway and condition in Supplementary Material (Supplementary Tables ST9), and for discussion purposes, we present the bar chart for results found for parameters in the Notch pathway in Figures 8, 9. Figure 8 presents these effects for the normoxia condition with similar cell stimulation (Figure 8A) or different cell stimulation (Figure 8B). Figure 9 shows the PRCC results for the hypoxia condition comparing normoxia vs. hypoxia (left panel) and 1 h vs. 12 h under hypoxia (right panel).

**FIGURE 8.**
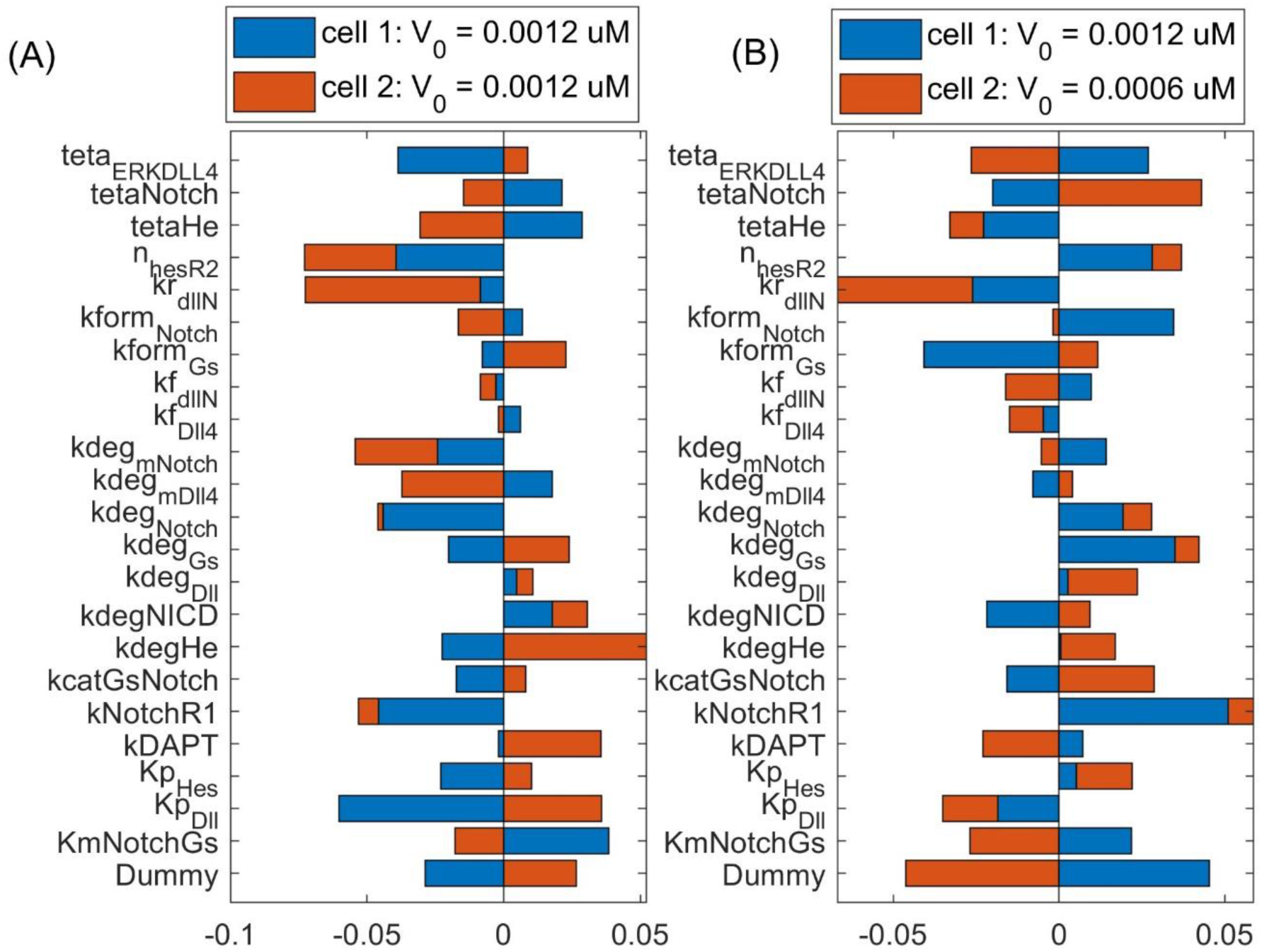
Sensitivity analysis of Endothelial Cell Pattern Index (ECPI) of each cell to changes in the parameters (in the Notch pathway) under normoxia considering (A) the same initial VEGF stimulation for the two cells or (B) different initial stimulation for each cell.

**FIGURE 9.**
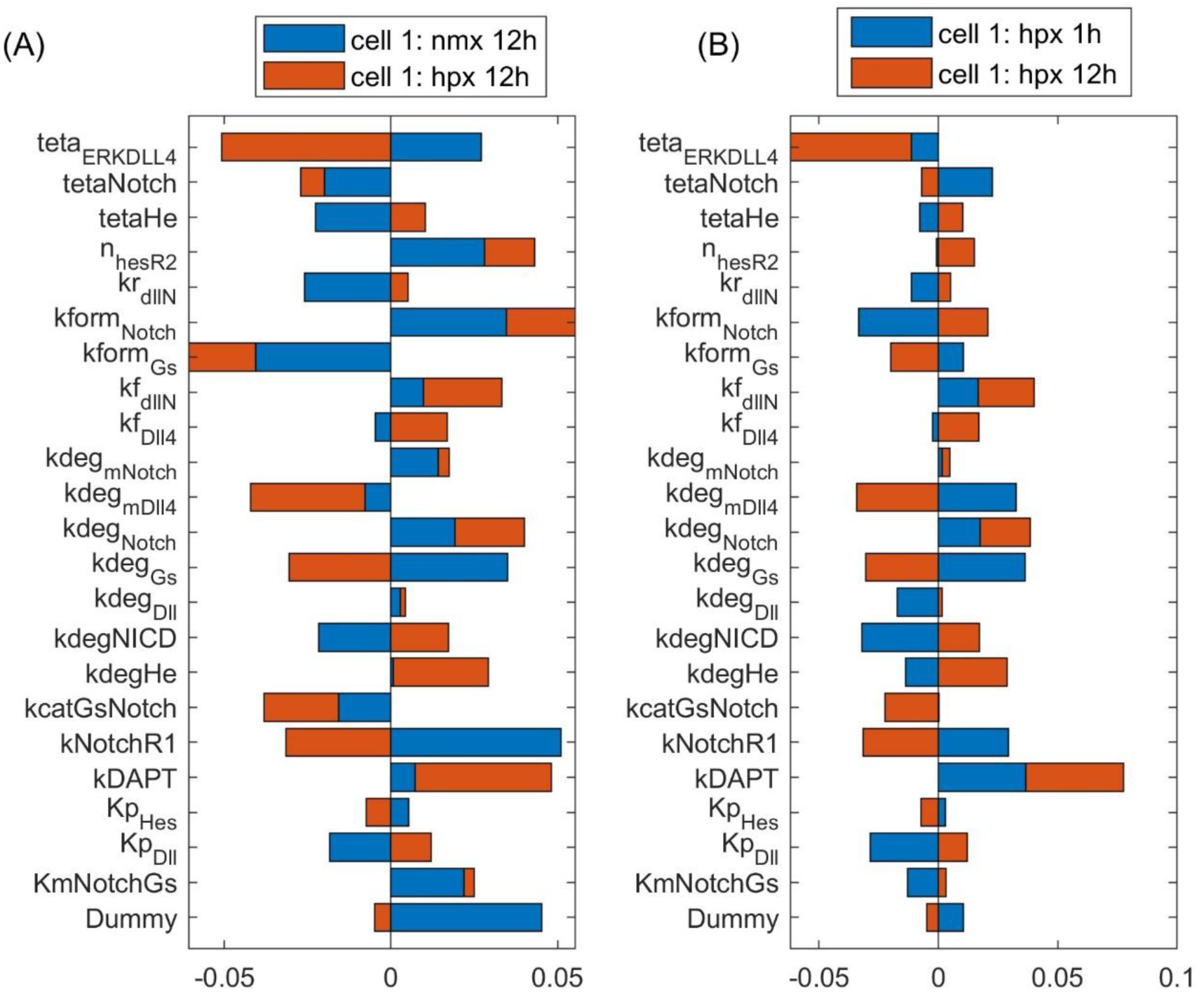
The influence of parameters on the ECPI is altered by oxygen availability (A) and time under hypoxia (B). Results shown for the first cell with differential cell stimulation (VEGF at cell 1 = 2* VEGF at cell 2), for parameters in the Notch signaling pathway.

Observing the influence of parameters from the different pathways, we initially note that regardless of the VEGF stimulation of the two cell (whether under the same or different stimulation), the parameters that mostly influence the ECPI of the second cell remain the same (degradation of VEGFR2 bound to NRP1—kdegR2NRPnp, phosphorylation of Raf— kmPKCRaf, formation of PIP2—kgenPIP2, dephosphorylation of sphingosine—kdpS1P, upregulation of sphingosine—kcatSK1Sph, phosphorylation of Raf—kcatPKC, and activation of IP3—KmPIP2PLCy). For the first cell, on the other hand, the parameter’s influence is more distributed among parameters. Among parameters in the Notch pathway, the degradation rate of Hes1 (kdeg_He) is the most influential for the second cell, and the Michaelis-Menten constant for the production of Dll4 (kp_Dll) is the most influential for the first cell. The Hill coefficient for VEGFR2 repression by Hes1 (n_HesR2) imposes a higher influence on the second cell ECPI but also affects the first cell. Interestingly, under differential stimulation of the cells, n_HesR2 has a positive PRCC value, as opposed to the negative PRCC seen with cells under the same stimulation (Figure 8A), and a similar effect is seen for kNotchR1 (the multiplication factor for the upregulation of VEGFR1 due to Notch pathway stimulation). This corroborates the notion that patterning of the cells is dependent on VEGF stimulation, and differential stimulation affects how the cells respond.

Comparing the sensitivity analysis of the ECPI of the more stimulated cell (cell 1) under hypoxia (12 h), the most influential parameters (PRCC >0.05) on the VEGF pathway are kNRP1VEGFR2off, khesr2, kp_iVR2, kvr1on,krec_iR2, and kvr1off. In the Notch pathway, we have teta_ERKDll4, kform_Notch, and kform_Gs. In the NO pathway, kr_paktHsp, and khif1eNOS. In the HIF pathway, ktranslmHIF1a, kdeg_HIF1a, and k12_degHIF1a. In the Calcium pathway, we have kon_DAGPKC, koff_capkc, kf_CaCaM, kcatERK, and Caext. Finally, in the ERK/Akt pathways, the most influential parameters are kr_AktPIP3, koffSK1, kformAkt, kdp_Akt, kcatSK1Sph, and kRasGAP. Comparing the number of influential parameters in each pathway, we note the parameters in reactions in the VEGF and ERK/Akt pathways. Regarding the power of influence (higher PRCC), the most influential parameters are in the VEGF and ERK/Akt pathways.

To evaluate the effects of hypoxia on parameter influence, we compare the PRCC of the most influential parameters under normoxia (12 h) and hypoxia (12 h) on the more stimulated cell. For parameters in the VEGF and HIF pathways, we note that hypoxia increases the influence of these parameters on defining the PRCC. This effect is also seen for some parameters in other pathways. However, in several cases, for instance, teta_ERKDLL4, kdeg_mVEGFA, kNotchR1, delta_ox22, kdpAkt, koffSK1, and Caext, hypoxia reverses the effect of the parameters (+ PRCC to—PRCC or vice-versa). Our results indicate that hypoxia is an effector of EC patterning guidance, by modulating how parameters influence cell patterning under differential VEGF stimulation. Following up on this finding, we also investigated the effect of time under hypoxia, comparing the influence of parameters on defining the ECPI of cell 1 under 1 h or 12 h of hypoxia conditions, with differential cell stimulation (VEGF_c1 = 2*VEGF_c2). The results indicate that, for the most influential parameters, a longer time under hypoxia conditions increases the effect of the parameters seen at 1 h of simulation. However, for some of the most influential parameters (e.g., O_2_, k12_degHIF1, Km_IP3R, kp_pAxl, and kf_eNOSHSP), time under hypoxia reverses the effect of the parameters on determining the ECPI. In summary, our sensitivity analysis under hypoxia indicates that oxygen levels and time under hypoxia influence cell patterning and should be taken into consideration for pattern control strategies.

### 3.3 Differential exogenous VEGFA stimulation is required for cell patterning

To observe cell patterning under different VEGFA stimulation for normoxia, we simulate the time-courses of VEGFR2 (Figures 10A), pVEGFR2 (Figures 10B), Hes1 (Figures 10C), Dll4 (Figures 10D), and NICD (Figures 10E) under normoxia, with one cell being more stimulated (VEGFA = 50 ng/mL) than the other (VEGFA = 25 ng/mL).

**FIGURE 10.**
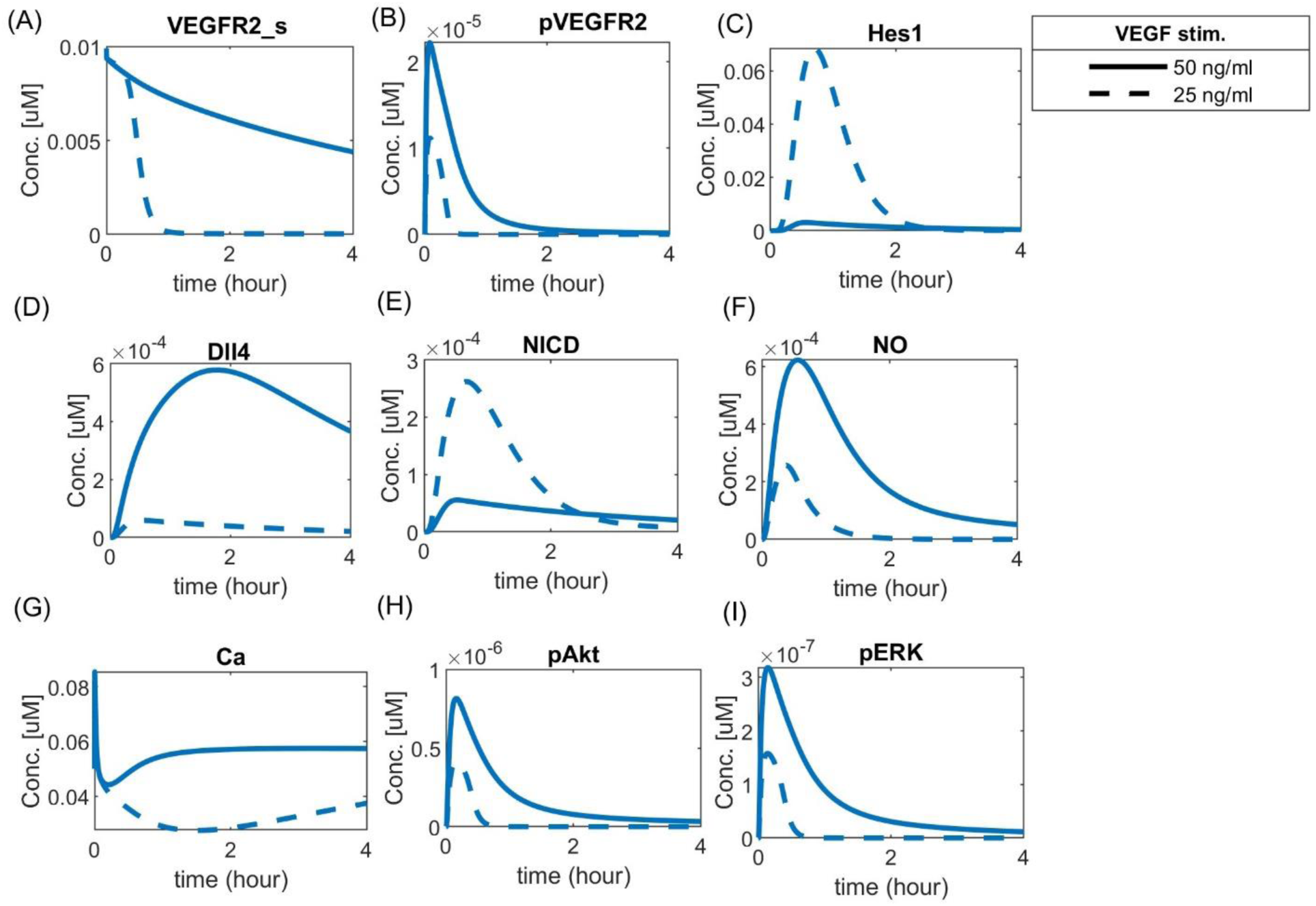
Cell patterning under normoxia with VEGF stimulation. Each cell (solid or dashed lines) was stimulated with a different amount of VEGFA, leading to a clear patterning response guided by the Notch pathway. Predicted responses of (A) Surface VEGFR2, (B) Phosphorylated VEGFR2, (C) Hes1, (D) Dll4 protein, (E) NICD, (F) Nitric Oxide (NO), (G) Calcium++ (Ca), (H) Phosphorylated Akt, and (I) Phosphorylated ERK1/2.

The two cells present different concentration dynamics over time of the analyzed species. The more stimulated cell presents increased levels of surface VEGFR2, phosphorylated VEGFR2, and Dll4, while the unstimulated cell presents higher NICD and Hes1 levels, consistent with the expected behavior of tip and stalk endothelial cells (Venkatraman et al., 2016; Akil et al., 2021). The effect of Hes1 in inhibiting VEGFR2 is shown by the fast decay seen as Hes1 increases in the less-stimulated cell. Additionally, the combined effects of VEGFA and Notch signaling are noted downstream, with a higher amount of NO, Ca++, pAkt and pERK1/2 in the more stimulated cell than in the less stimulated cell (Figures 10F–I, respectively).

To compare the effects of hypoxia with the noticed behavior, we simulate the time-courses for hypoxia without exogenous VEGFA and with exogenous differential VEGFA (Vext = 0.00012 μM) to that of normoxia with both cells receiving a similar basal stimulation (VEGFA = 0.000001 μM) (Figure 11). In this condition, we can see that hypoxia upregulates the simulated species compared to normoxia, but no patterning occurs unless exogenous VEGFA is added to promote a differential stimulation of the cells. This complies with the assumption used in our model that the autocrine VEGFA does not lead to patterning, and an exogenous differential stimulation must be present for patterning to occur. This assumption is based on previous studies comparing autocrine and paracrine VEGF signaling in endothelial cells, as discussed in the methodology (Lee et al., 2007). As an additional analysis, we investigate the effect of inhibiting the Notch pathway using the *γ_secretase_* inhibitor DAPT (N-[N-(3, 5-difluorophenacetyl)-l-alanyl]-s-phenylglycinet-butyl ester)) (Supplementary Figure S14). Our simulations show that DAPT [20 μM] is able to inhibit VEGF-induced NICD and Hes1 expression, leading both cells to present a similar pattern and expression profile, reproducing results seen *in vitro* (Takeshita et al., 2007).

**FIGURE 11.**
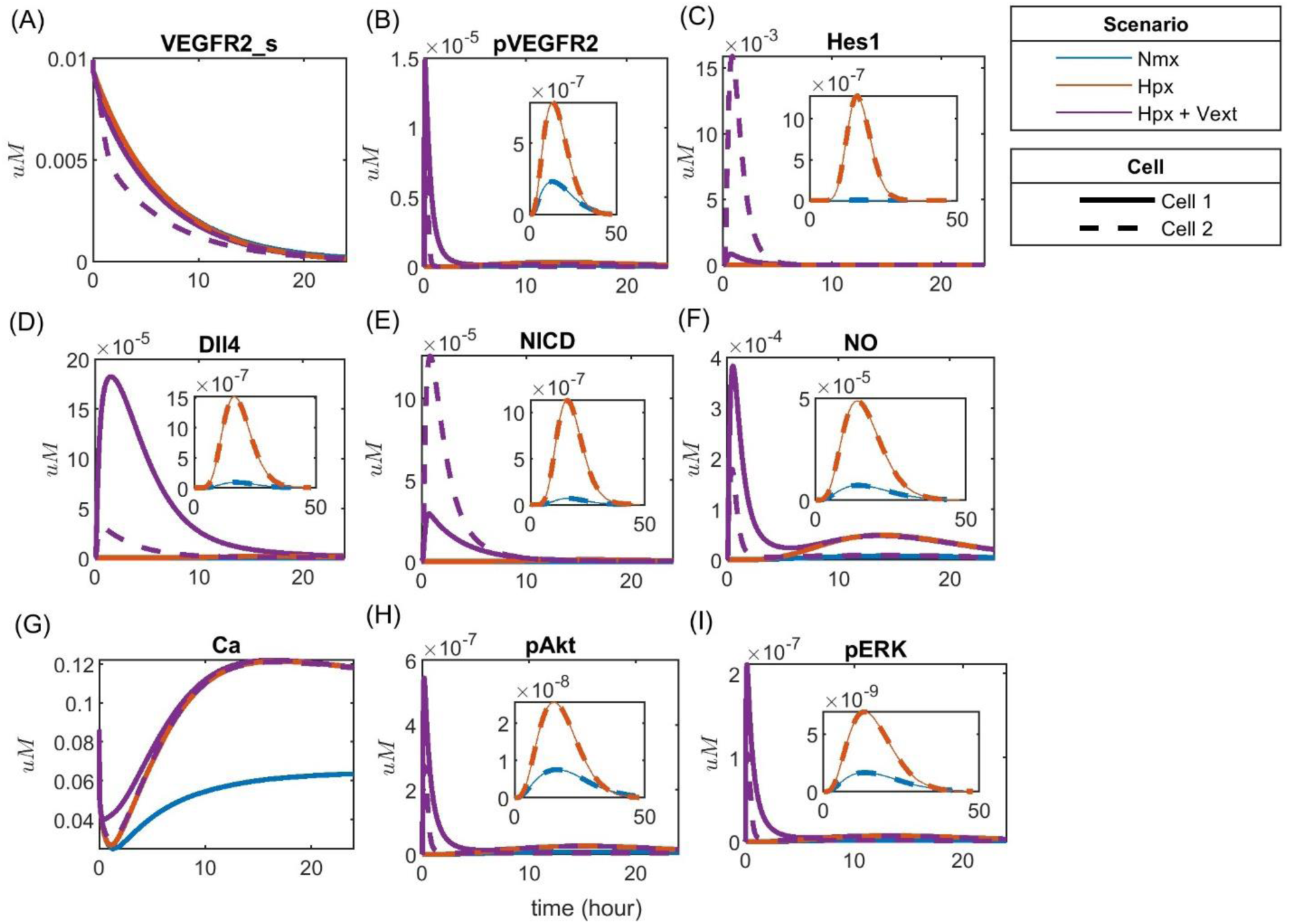
Patterning under hypoxia with and without external VEGFA stimulation for (A) Surface VEGFR2, (B) Phosphorylated VEGFR2, (C) Hes1, (D) Dll4 protein, (E) NICD, (F) Nitric Oxide (NO), (G) Calcium++ (Ca), (H) Phosphorylated Akt, and (I) Phosphorylated ERK1/2.

### 3.4 Notch signaling modulates vascular permeability through VEGF-induced NO expression

Given the known effect of the increase in vascular permeability due to VEGFA, we then investigated the mechanistic effect of Notch signaling on vascular permeability. Although initially controversial, studies performed in eNOS knockout mice showed that NO derived from eNOS under VEGF stimulation causes hyperpermeability (Duran et al., 2010). The leaky vasculature is also one of the issues noticed in therapeutic angiogenesis treating PAD with VEGFA (Han et al., 2022). To investigate the effect of Notch signaling on NO expression under hypoxia, representing the PAD condition, we simulate the time course of phosphorylated eNOS and NO in the two cells, at three different concentrations of Dll4 (0 μM, 2 μM, and 20 μM). Our results are presented in Figure 12. The simulations indicate that activation of the Notch pathway through Dll4 in one of the cells (in the simulation, cell 2) affects NO release by the 2 cells (panel A). In this simulation, both cells are stimulated with similar amounts of VEGF (50 ng/mL). When both cells start with similar amounts of Dll4 (blue lines), NO release is similar. However, as we alter Dll4 in the second cell (increasing the activation of the Notch pathway), the response differs. Observing the response to 0.001 μM of Dll4 in the second cell (green lines), we note that both cells are affected.

**FIGURE 12.**
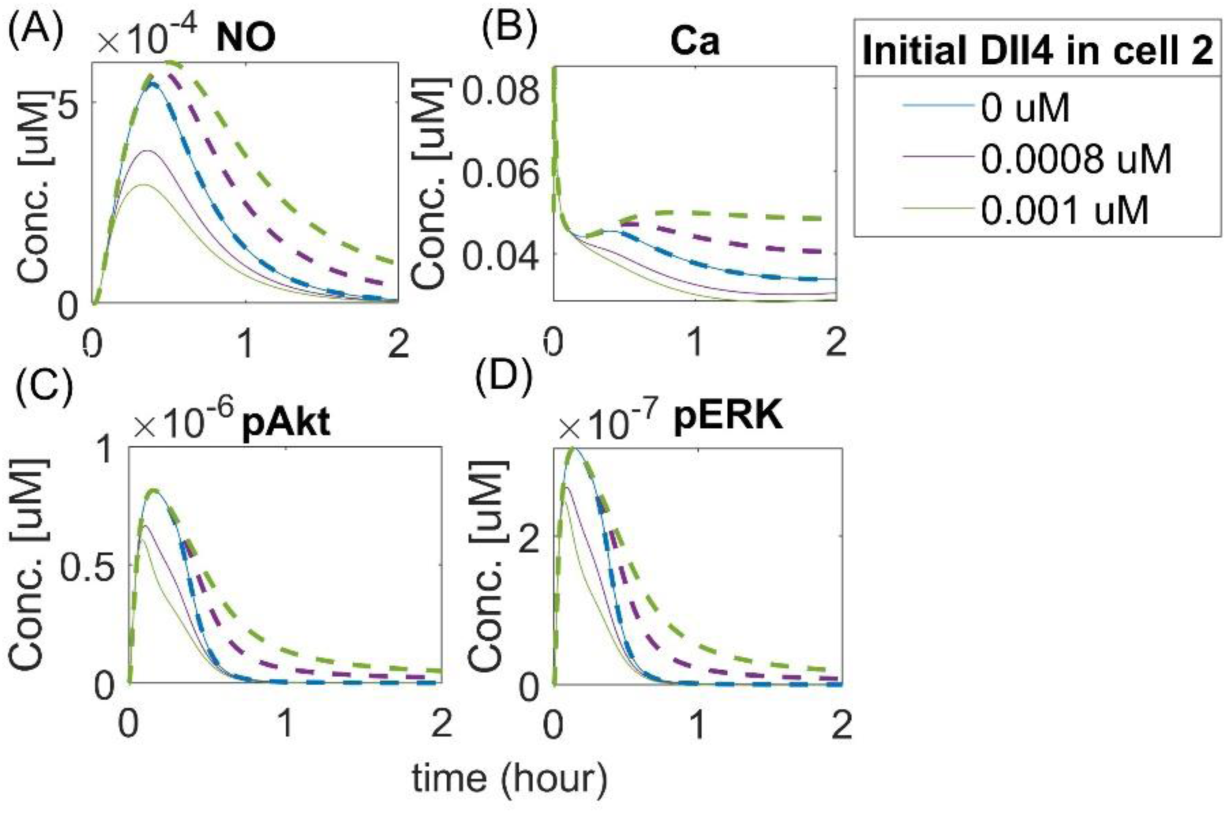
Activation of the Notch pathway regulates NO release differentially in neighbor EC. Predicted responses of (A) Nitric Oxide (NO), (B) Calcium++ (Ca), (C) Phosphorylated Akt, and (D) Phosphorylated ERK1/2. The dashed line represents the response of cell 2, and the solid line represents the response of cell 1. Different colors represent different initial concentrations of Dll4 in the second cell.

Our results indicate that NO release by the first cell is smaller than its release by the second cell. Increasing Dll4 in the second cell activates Notch signaling in the first cell, leading the first cell to assume the stalk position. The first cell then has the amount of free VEGFR2 limited, which leads to less activation by VEGF and reduced NO release (as a downstream effect of VEGFR2 activation). It is interesting, however, that we also see an increase in the release of NO by the second cell (which, in this case, assumes the tip position, compared to the 0 initial Dll4 condition (blue lines). We note a similar effect of intracellular Ca++ dynamics (panel B). Evaluating the effect on Akt and ERK phosphorylation (panels C and D), we also note an initial difference (comparing the blue line with purple and green lines), but with a faster co-incidence of the time-course behavior for these species (the solid green and purple lines representing the second cell, which acts as the tip cell, merge with the blue line before 1 h of simulation) compared to Ca++ and NO (where the merging occurs only after 1 h for NO, and does not merge with the blue line during the 2 h simulation for the Ca++ simulation). These results indicate that Notch activation by Dll4 upregulation in one of the cells affects downstream signals in both cells, including NO release, which indicates that vascular permeability might be affected by Notch activation, with tip and stalk cells presenting differential NO release due to Notch activation.

## 4 DISCUSSION

Sprouting angiogenesis as a response to hypoxia is a necessary process tightly regulated by an amalgamate of agents and signaling pathways (Hashimoto and Shibasaki, 2015; Naito et al., 2020; Rodriguez et al., 2021; Zhang et al., 2022). Endothelial cell patterning into the tip or stalk phenotypes guide sprout formation, promoting blood flow restoration to hypoxic tissues, such as seen in the context of diseases like PAD (Gustafsson et al., 2005; Annex and Cooke, 2021; Han et al., 2022). Efforts to characterize, understand, predict, and control cell behavior under abnormal conditions are relevant in this field, as they can help develop new therapeutic strategies and interventions in angiogenesis.

In this study, we propose a structured, state-of-the-art methodology to comply with good modeling practices and reduce the computational cost required for fitting and simulating large models. We present the step-by-step procedure to perform each of the main parts of model design, training, and validation, and we apply this methodology to a new integrative model of the main pathways in EC during angiogenesis. Through this kinetics-based model, we represent and simulate the interaction and patterning of 2 ECs stimulated with VEGFA under normoxia and hypoxia conditions. This model was developed as a first step towards investigating the differential signaling and response of each of the two cells, as they are differently stimulated by VEGFA and interact through Notch signaling under varying oxygen conditions. We calibrated the model using data from experiments performed in endothelial cells for both oxygen conditions and defined a ratio for stalk/tip pattern identification based on known ligands and receptors overexpressed in each of these phenotypes (del Toro et al., 2010; Xu and Li, 2022). Considering recent efforts in designing and characterizing “virtual cells” (Zhao et al., 2021; Lim et al., 2022; Zhang et al., 2022), this work contributes to the field by bringing insights into cell-cell interaction, patterning, and signaling with the potential translation of results to pathological conditions (specifically, hypoxia-characterized diseases such as PAD).

As new experimental results become available, revisiting previous assumptions and models is essential, making adjustments that bring us closer to *in vivo* conditions. Endothelial cell patterning and interaction is a significant field of study in therapeutic angiogenesis since these cells are important determinants of angiogenesis and blood flow restoration (Qiu and Hirschi, 2019; Lim et al., 2022). Considering the many aspects that influence their heterogeneity (Gifre-Renom et al., 2022), a computational approach provides the flexibility required to implement further changes and test hypotheses. Our model reproduces the characteristic EC patterning behavior under normoxia given differential VEGF stimulation (Figure 10) and predicts a distinct behavior under hypoxia (Figure 11). It also supports the requirement of a paracrine differential VEGF stimulation for cell patterning. Additionally, it indicates differential signaling and patterning under hypoxia between the two cells, which is supported by previous studies showing that hypoxia regulates and promotes tip/stalk differentiation through overexpression of Dll4, indirect regulation of VEGFR2 and NRP1 established by Notch signaling, and upregulation of Hes1 (Carmeliet et al., 2009; Rodriguez et al., 2021).

Tip and stalk endothelial cells interact and respond differently to signals and targeting them has a therapeutic potential in terms of tip-stalk cell control and selection (Chen et al., 2019). Through global sensitivity analysis (Figures 8, 9; Supplementary Table ST9), our model allowed us to investigate this distinct behavior in a system of two interacting cells, focusing on which parameters and reactions are more influential in defining each cell’s pattern. Although we expected parameters from the Notch pathway to be the most influential in determining cell patterning, our simulations indicate that, although they play a part in modulating the ECPI, parameters in reactions involved in the VEGF, ERK, and Akt pathways have a higher influence on the ECPI. As these pathways’ activation leads to Notch signaling activation, their influence on the ECPI becomes clear in an integrative model such as the one we present. ECs require differential VEGF stimulation to assume a pattern (tip or stalk), and similar stimulation turns cells toward an undifferentiated state. Previously, others have reported that EC can assume tip, stalk, and quiescent phalanx states, where the latter refers to more mature EC, formed once the vessel is perfused (Pasut et al., 2021). Partial active/inactive states during differentiation have also been reported through computational models (Venkatraman et al., 2016). These two states are seen previous to the point where the cell assumes the tip or stalk position, under stimulation by VEGF. In this work by Venkatraman et al., EC quiescence is noted for similar VEGF stimulation (VEGF at cell 1 = VEGF at cell 2 = 0) and led us to question if the parameters that define cell patterning differ and, if so, how they differ when cells are differentially or similarly stimulated or under different oxygen conditions. Through global sensitivity analysis of the ECPI of both cells subject to similar (Figure 8A) or different (Figure 8B) VEGF stimulation, we found that for several parameters, the differential stimulation that guides patterning leads to a change in parameter effect (moving from negative PRCC to positive PRCC or vice-versa), but this effect was parameter-dependent. This simulation advocates for cell patterning as being a highly regulated event that depends on the intensity of VEGF signaling being sensed by each cell. Our model is also able to reproduce the effects of DAPT inhibition of the Notch-induced cell patterning post VEGF stimulation (Supplementary Figure S14) seen experimentally (Takeshita et al., 2007).

In this work, we assume that hypoxia’s effects on driving angiogenesis are mostly due to the paracrine effect of VEGF released by other cells on endothelial cells. This is based on previous evidence, where the autocrine effect of VEGF (generated by EC under hypoxia) is shown to be mostly for cell survival, with few contributions to angiogenesis (Lee et al., 2007). However, recent work indicates otherwise (Jin et al., 2019), with VEGF production by EC under hypoxia leading to angiogenesis. We hypothesize that this difference is due to the cell environment tested in each experiment, as exogenous VEGF stimulation by other cell types might shadow the angiogenic effects of autocrine VEGF. Additionally, we assume that the effects of hypoxia on regulating VEGF receptors on cell surface depend on Notch signaling, which we pose as a question to be answered in future experiments. Future experiments should also assess the extent to which Notch signaling participates in defining vascular permeability through NO regulation and whether this interaction interferes with vascular permeability and leakiness profile in pathological conditions such as PAD. Many methods are available to estimate NO *in vivo* and *in vitro* (Goshi et al., 2019), and previous works have estimated NO release by HUVECs to be within the ranges of nM to μM (Østergaard et al., 2007; Ugusman et al., 2014; Janaszak-Jasiecka et al., 2018). Additionally, *in vivo* measurements of NO report values of about 1 μM, with the values of EC50 for soluble guanylate cyclase (sGC) in the vascular smooth muscle as low as several nM (Chen et al., 2008). Our simulations predict concentrations in the order of 10^−1^μM. This difference can be due to the initial concentration of NO assumed (0 μM), and more precision of absolute values estimates might require fitting the model to absolute values of NO, instead of normalized to their maximum concentration only. This assessment should also be included in future works. Despite this difference in the order of magnitude predicted by our simulations, compared to experimental *in vitro* data, our model qualitatively indicates the difference in NO production by the two cells, which was among our stated goals.

The methodological approach we used in this work allowed us to develop the model and perform simulations within reasonable computation time, despite the large size of the model. The modular implementation during global optimization aided in speeding up the process of model fitting. Performing SIA and PIA as a way of achieving good modeling practices and limiting the number of parameters for fitting is an additional aspect that helped in this regard. Our method is, therefore, efficient for working with larger mechanistic computational models. Recently, a new framework and steps to build large mechanistic models was proposed, integrating annotated input text files for specific data, python-based platforms for processing the input files and generating Antimony files to be converted to SBML standards and Python for model simulations (Erdem et al., 2022). Another computational framework for parameterization of large-scale mechanistic models has been proposed, with significant advancement on computation time for very large models (>1000 parameters) (Fröhlich et al., 2018). In this framework, the authors include practical identifiability analysis not as a step prior to optimization/fitting, but as an investigation of prediction uncertainty caused by parameter uncertainties performed post-fitting. Their results pointed to most of the calibrated parameters being poorly identifiable, but still allowing them to obtain low-uncertainty predictions. Practical identifiability requires both a sensitivity analysis (to verify that the parameters are influential to the observables in question) and a collinearity analysis (to exclude collinear parameters), and performing these evaluations before the final optimization of the values helps limit the number of parameters being fitted (therefore, the computational cost and time of model optimization) and improve prediction uncertainty caused by parameter uncertainty. Although the method we proposed in this work relies on a simple collinearity analysis (removing parameters with opposing effects before fitting), this method could be automatized by using platforms such as VisID (Gábor et al., 2017). An improved approach that could be tested in future works is to perform the structural identifiability, followed by our simplified version of the practical identifiability (based on initial guesses for parameters within reasonable ranges), then the global optimization, and as an additional analysis to UQ, a new round of practical identifiability (performed with an automatic platform such as VisID). This method might reduce the number of unidentifiable parameters prior to fitting and provide a more accurate and trustable model.

Despite our efforts to calibrate the model with experimental data from the same line of endothelial cells, we were not always able to find the required data, considering the level of detail and ramifications of our model. Data from different cell lines might produce inaccurate simulations and results (Chi et al., 2003). Another limitation of our model is not including Jagged in the Notch signaling pathway, which should be accounted for in future studies, along with a more detailed model of the effects of hypoxia on the calcium signaling pathway (Berna et al., 2002) and of angiopoietin on the Notch signaling pathway (Machado et al., 2019). Our model is based on *in vitro* data for calibration and assumptions defined. Translating our results to *in vivo* conditions requires further experiments and mathematical and physiological assumptions not included in this work. Additionally, obtaining experimental data suggested by the present model would further validate the model. Our model is also limited by the pathways considered, as other signaling pathways might be affected by hypoxia and influence cell behavior, such as Ang-Tie, JAK/STAT, and TLR signaling. Those could be considered for future studies. We also note that in the model presented we consider that the effects of HIF1 and HIF2 are summative on inducing VEGFA signaling. Previous works have shown that these HIF1 and HIF2 undergo a switch regarding their time dynamics, to ensure a continuous activation of pathways response to hypoxia and prolong cell survival (Bartoszewski et al., 2019). However, others have shown that HIF1 and HIF2 differentially regulate the hypoxia response, in a context dependent fashion (Branco-Price et al., 2012). In fact, HIF2 in macrophages has been shown to induce an anti-angiogenesis response by inducing soluble VEGFR1, which sequesters free VEGFA from the environment (Eubank et al., 2011). In ECs, HIFs have also shown opposite effects on regulating interleukin 8 (IL-8), a promoter of angiogenesis (Florczyk et al., 2011). Although in out model such opposite effects are not included, the base model here provided can be further developed to represent and simulate additional differential effects of different HIFs.

Another limitation of the model is the estimation of parameters that have not been quantified in the literature or in previous models. Our simulations are constrained to the estimated values. As additional experimental data become available, this constraint could be revisited, and the estimates updated. Additionally, ECs under hypoxia act differently from ECs under hypoxia serum starvation (representing PAD conditions), which is a future aspect to be investigated with our modeling strategy. Previous works discuss the role of calcium oscillations on endothelial cell patterning (Debir et al., 2021). In this work, we consider the signaling through calcium in the endoplasmic reticulum and cytoplasm, with the extracellular calcium effect accounted for through CRAC channels calcium influx and the Ca++–Na + exchanger (under hypoxia). In future work, the focus could be more on the calcium effect on EC patterning, using a more comprehensive model of the signaling through calcium in EC under hypoxia.

Our modeling strategy in this work is non-spatial, differing from several other models in the field (Bentley et al., 2008; Reynolds et al., 2019), and focusing mostly on time-dynamics of species involved. Although observing spatial distribution of tip/stalk cells provides important insights on a tissue-level, for example, regarding cell distribution, diffusion of molecules and cell migration, obtaining a more focused view on intra and intercellular dynamics of a pathway as complex as the Notch pathway is also required to design and optimize pro-angiogenesis therapies focused on modulating Notch signaling. Non-spatial, ODE-based models are also more easily calibrated against experimental data, with clear modeling protocols. Given that non-spatial and spatial analysis are both integral parts for understanding the Notch pathway, its downstream effects and upstream regulators, a next step for this model is to integrate spatial analysis. This integration of ODE-based and spatial models has been recently shown on a viral infection and immune response model (Sego et al., 2021).

The Notch signaling has been a therapeutic focus for treatment of conditions related to angiogenesis. Notch inhibition has been applied to treating tumors (Akil et al., 2021; Jiang et al., 2022) and immune and inflammatory disorders (Rizzo and Ferrari, 2015; Allen and Maillard, 2021). Notch signaling is known to affect different cells, as previously discussed, and its therapeutic application requires a detailed understanding of its effects and its effectors. Our model provides a detailed integrative approach of evaluating Notch signaling and its effects in the context of angiogenesis, and can be extended to integrate Notch inhibitors (e.g., DAPT to simulate therapeutic applications of Notch in different pathological scenarios, including pro- and anti-angiogenic contexts (Niu et al., 2022; You et al., 2023), as represented in Supplementary Figure S14. We present this work as a first step in understanding the interconnection of pathways between endothelial cells during hypoxia-induced angiogenesis, as well as differences between tip and stalk ECs under different stimulatory conditions. We believe it expands the knowledge brought by previous studies in the field, and it is our first approach to working with larger, integrative models in ECs with the goal of building a network that can be used for therapeutic angiogenesis purposes. Additionally, it can be used to test hypotheses related to the pathways included (e.g., vascular permeability control through Notch signaling modulation). In summary, our model provides a highly integrative, data-driven platform that can be built on by simply adding pathway inhibitors or stimulators for testing angiogenesis-related therapies and to study differences in signaling between tip/stalk EC at varying oxygen conditions.

## DATA AVAILABILITY STATEMENT

The original contributions presented in the study are included in the article/Supplementary Material, further inquiries can be directed to the corresponding author.

## AUTHOR CONTRIBUTIONS

RO: Conceptualization, Data curation, Formal Analysis, Investigation, Methodology, Project administration, Software, Validation, Visualization, Writing–original draft, Writing–review and editing. BA: Funding acquisition, Supervision, Writing–review and editing. AP: Conceptualization, Formal Analysis, Funding acquisition, Investigation, Methodology, Resources, Supervision, Writing–original draft, Writing–review and editing.

## ACKNOWLEDGMENTS

The authors thank members of the Popel laboratory, Yu Zhang and Min Song, for helpful comments and discussions, especially on the model methodology.

