## Supplementary figures and images for "Endothelial cells signaling and patterning under hypoxia: a mechanistic integrative computational model including the Notch-Dll4 pathway"

### Figure S1.jpg

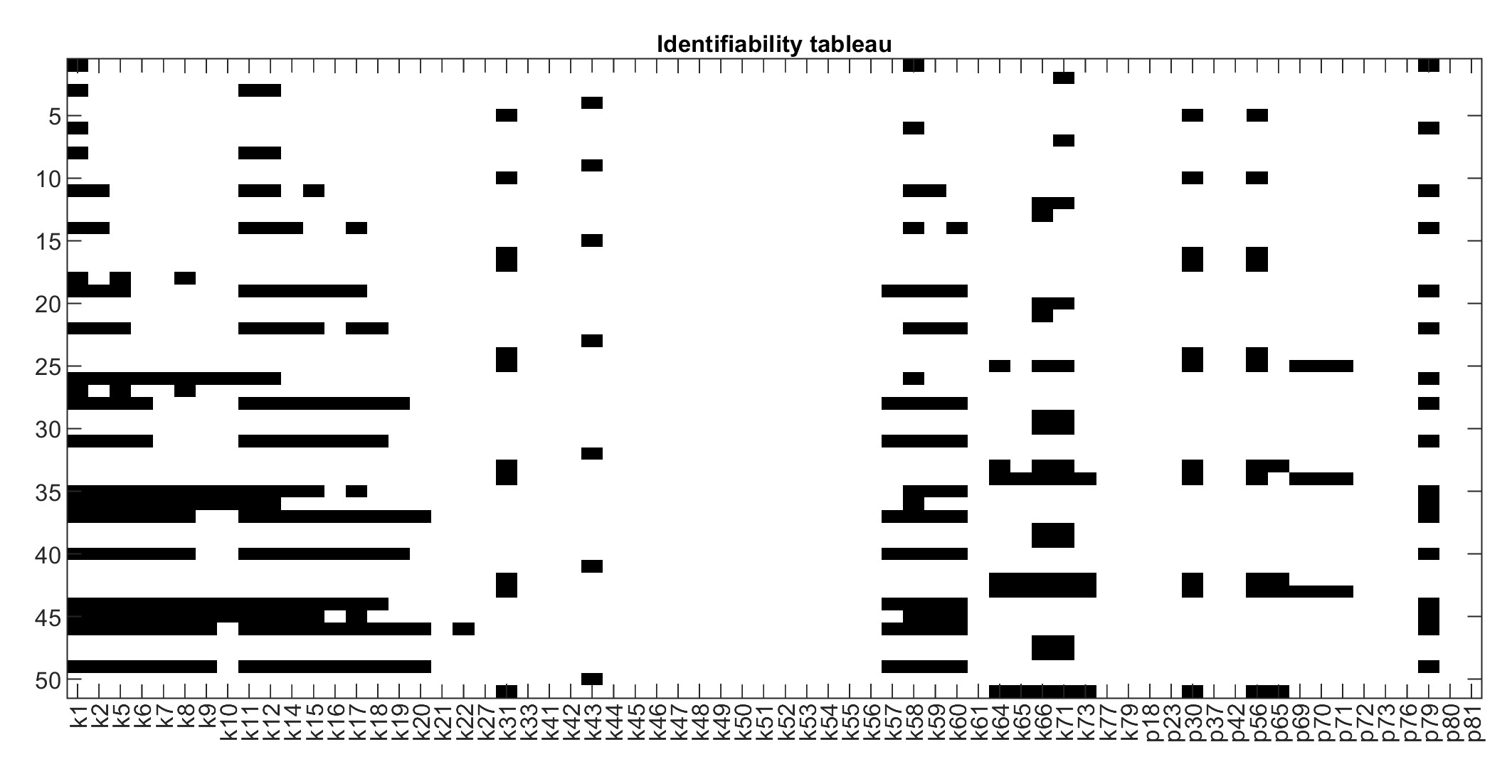

### Figure S2.jpg

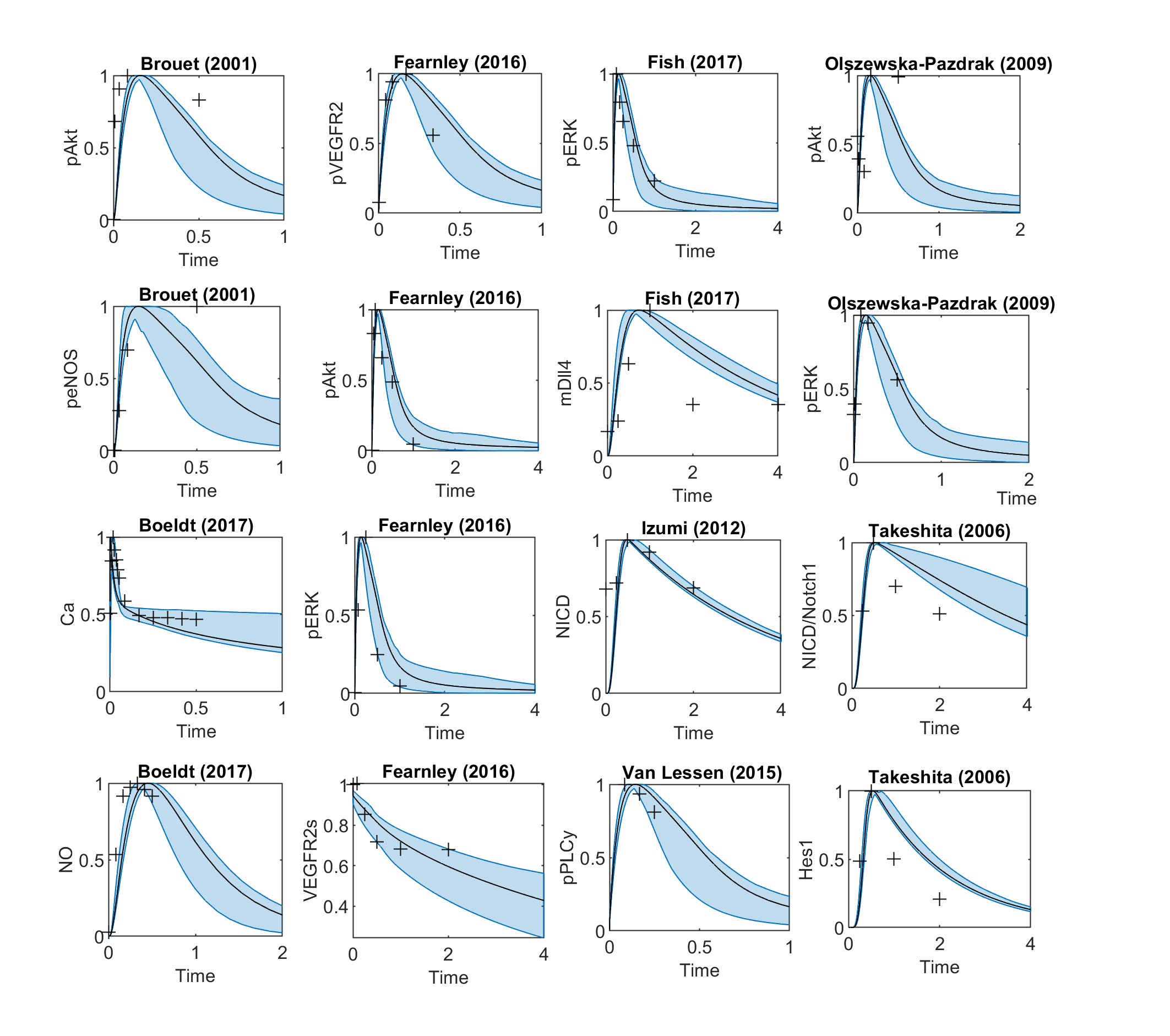

### Figure S3.jpg

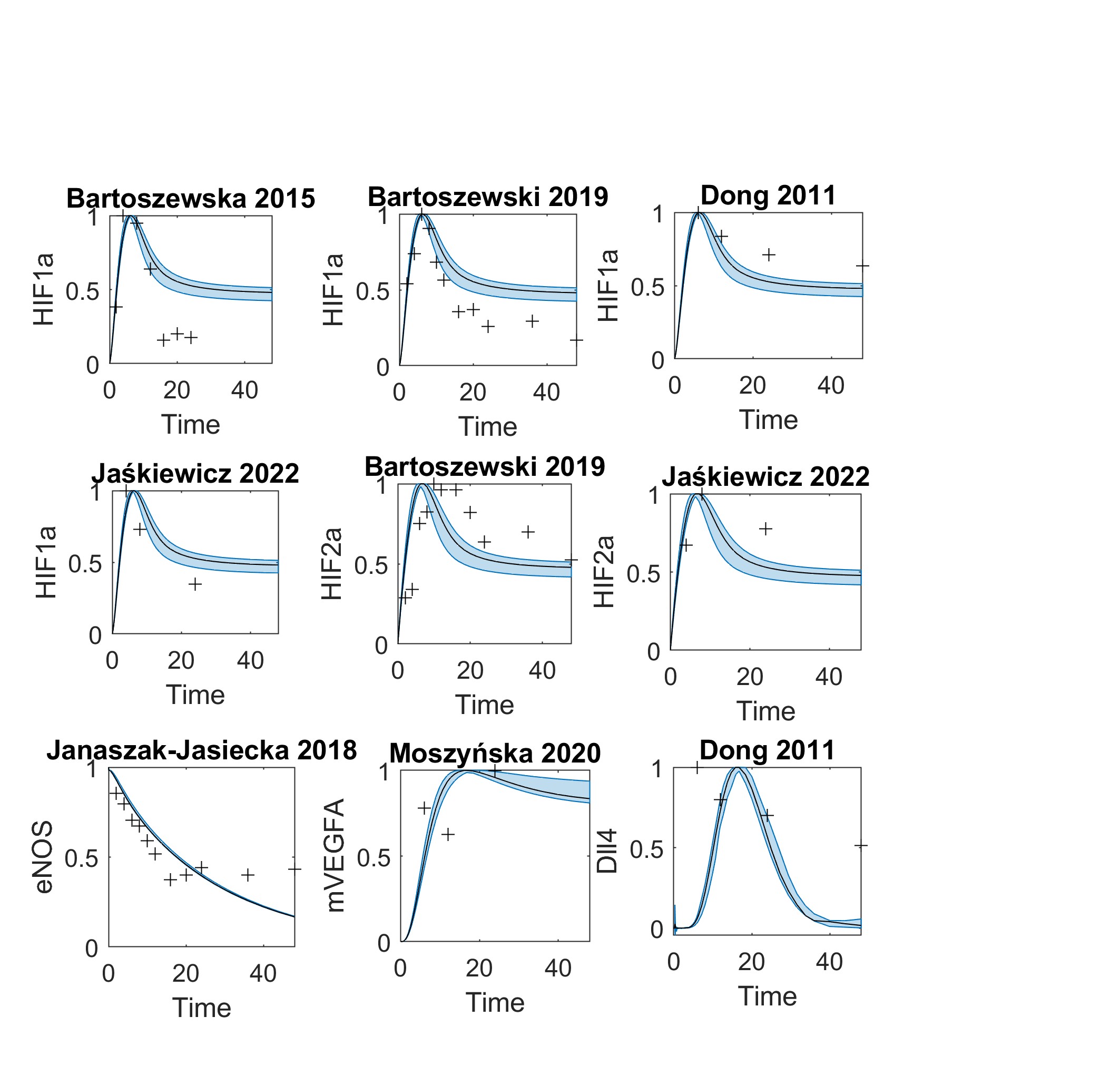

### Figure S4.jpg

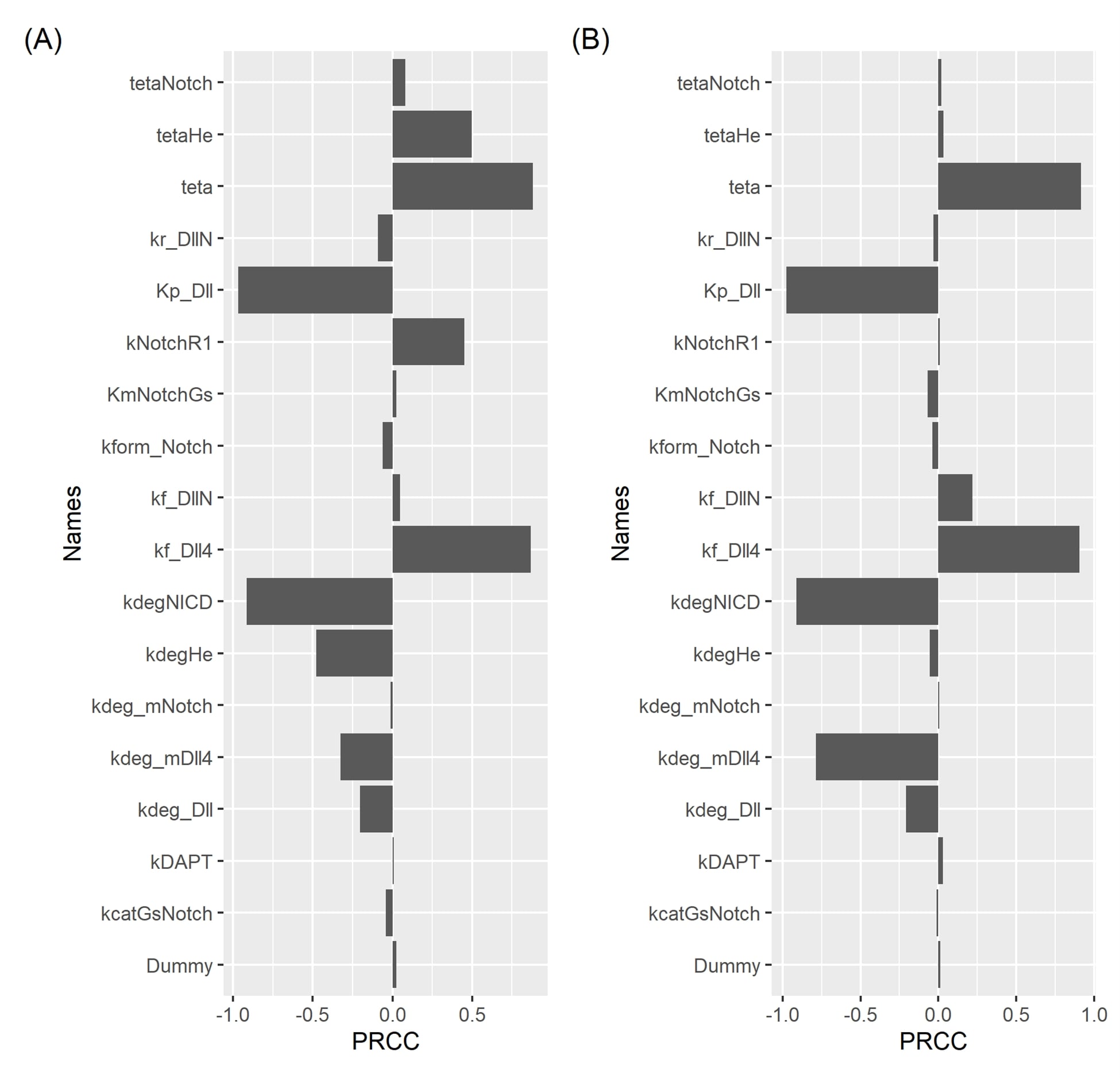

### Figure S5.jpg

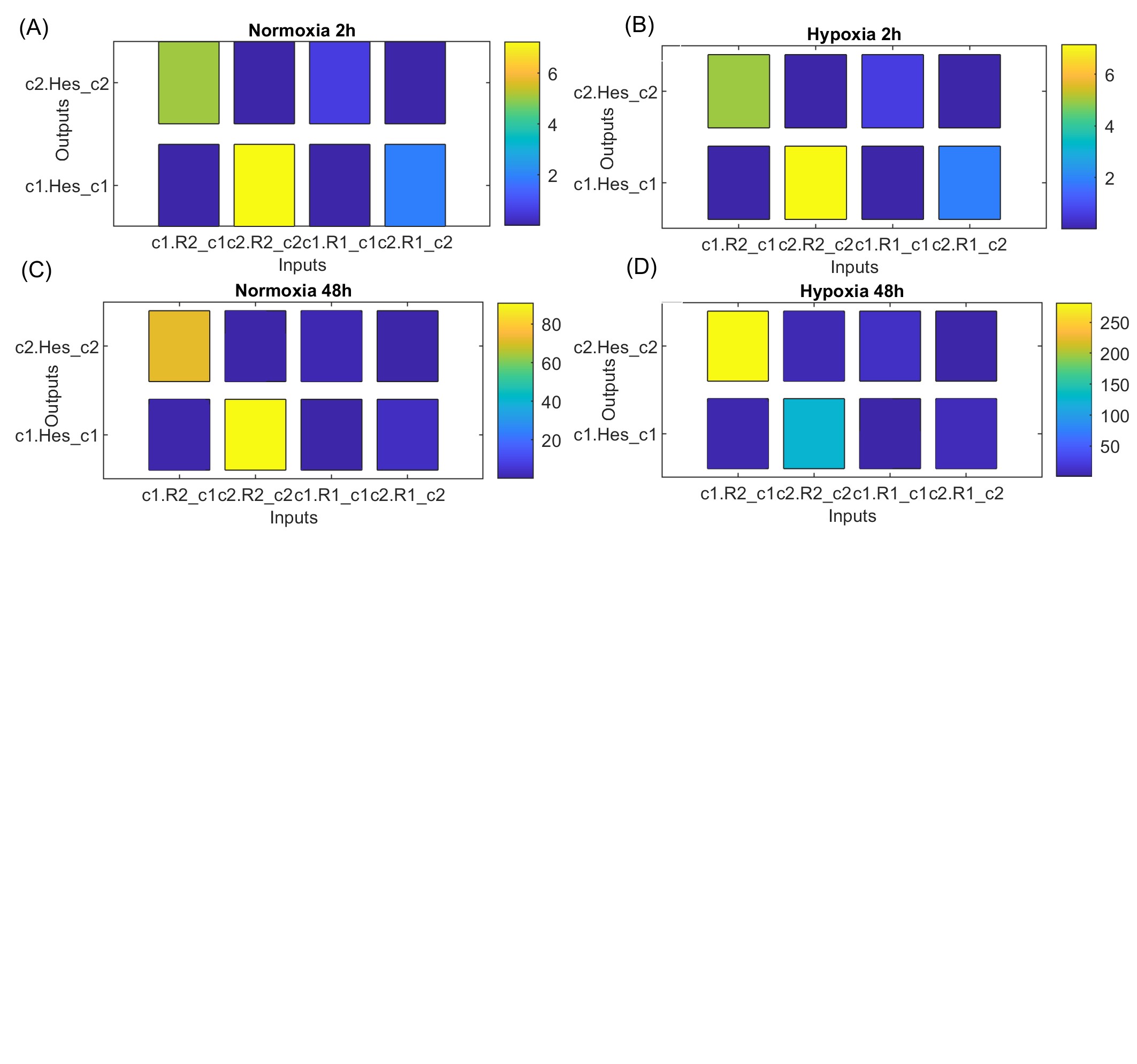

### Figure S6.jpg

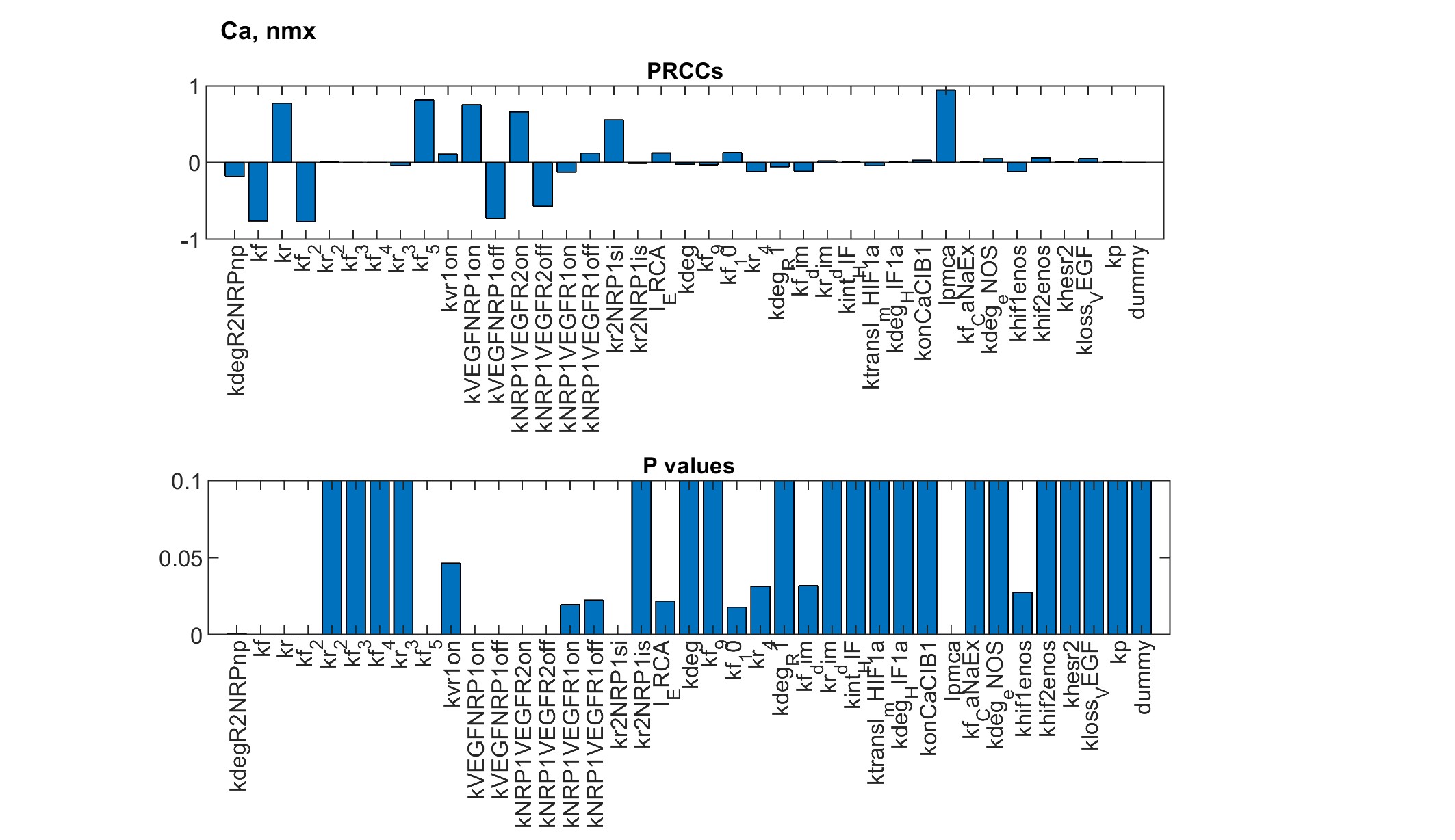

### Figure S7.jpg

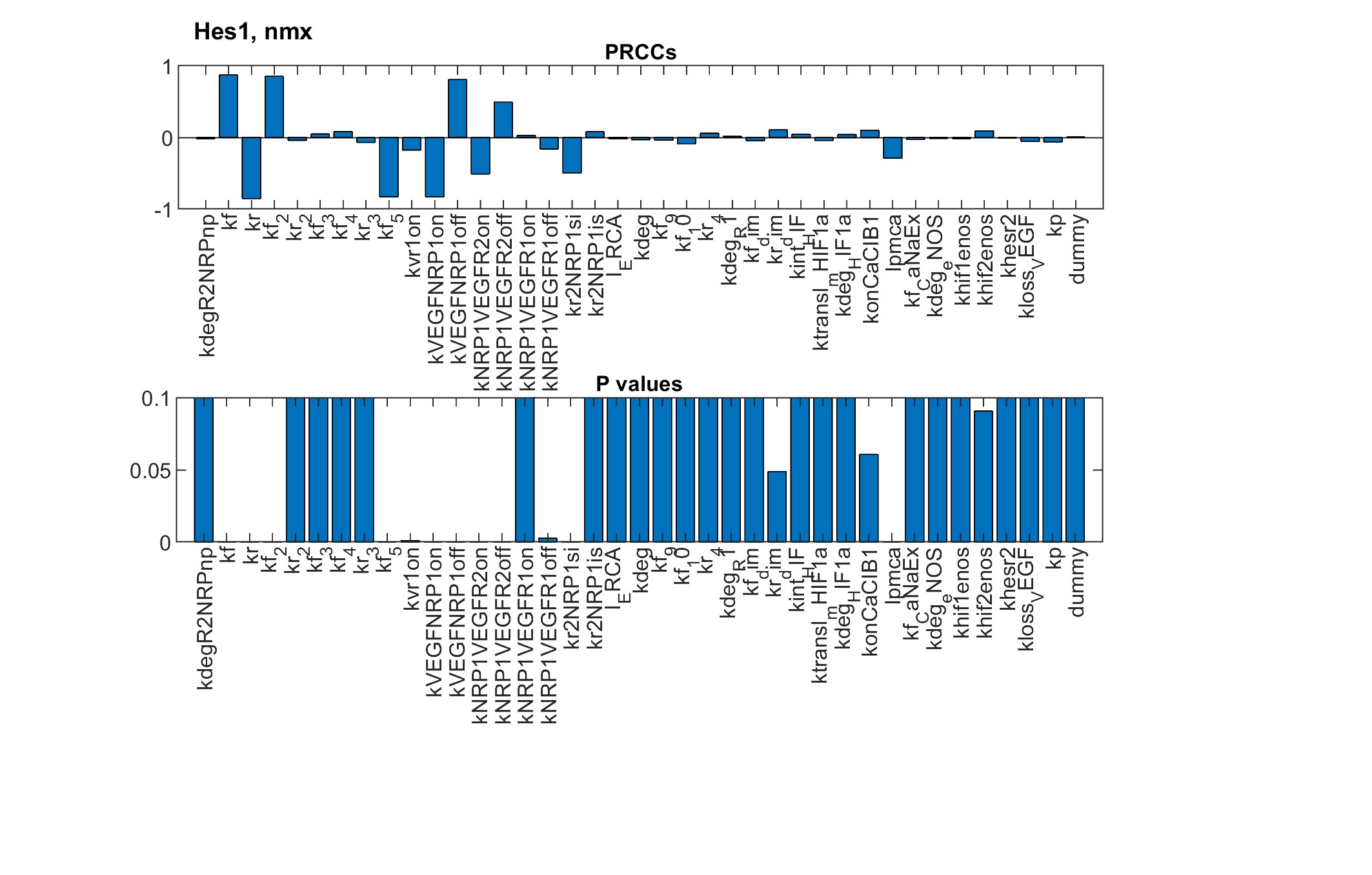

### Figure S8.jpg

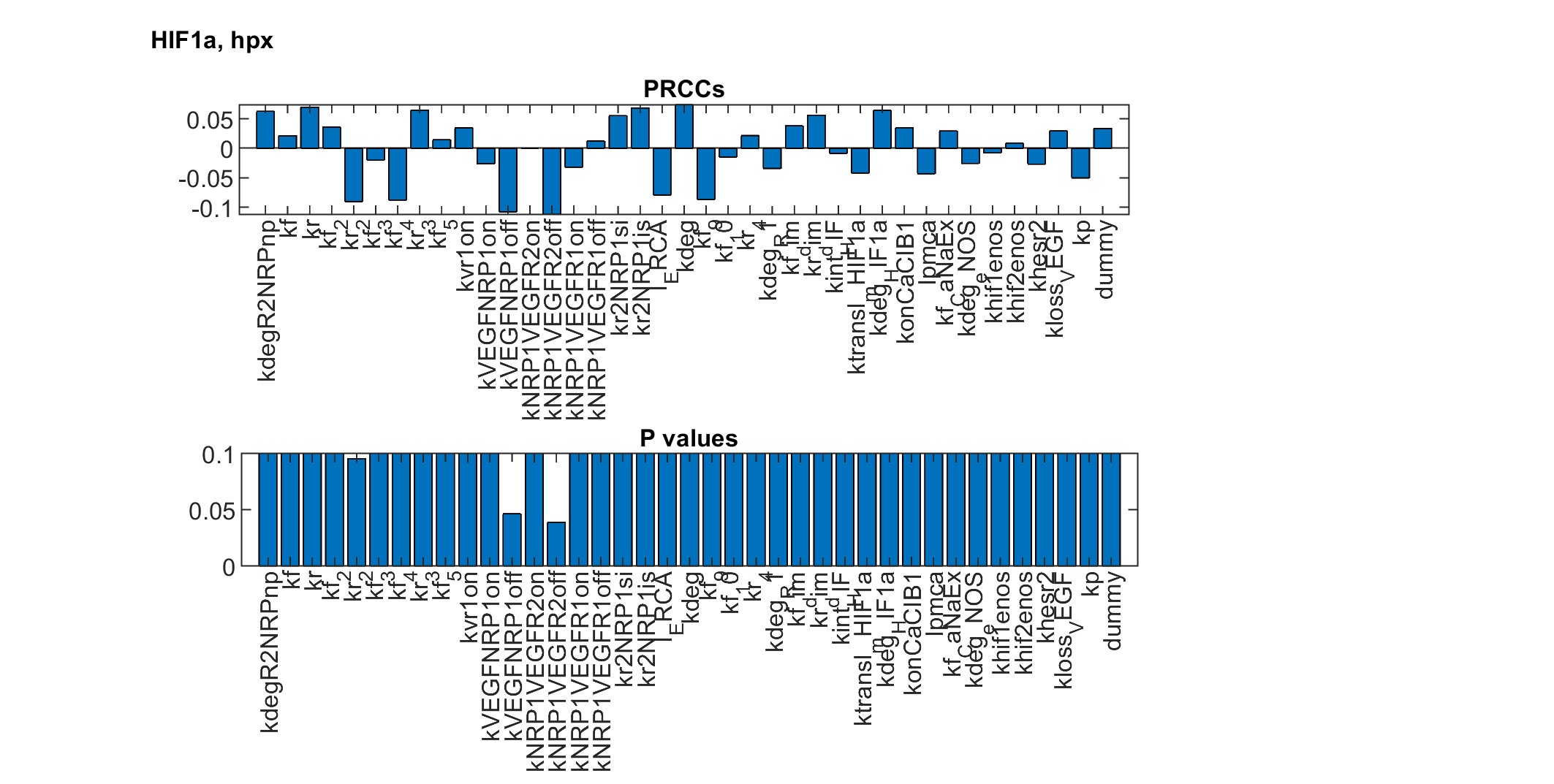

### Figure S9.jpg

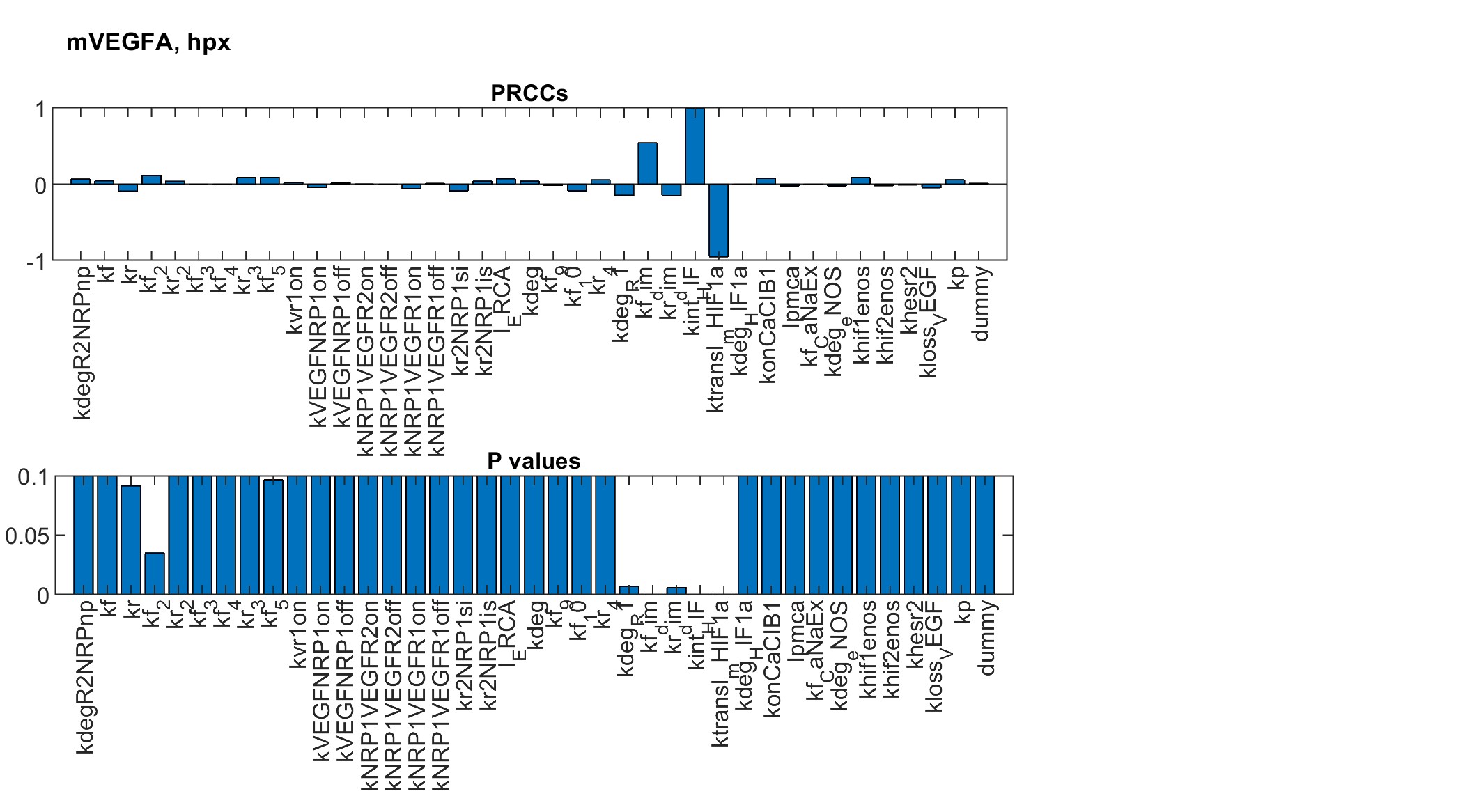

### Figure S10.jpg

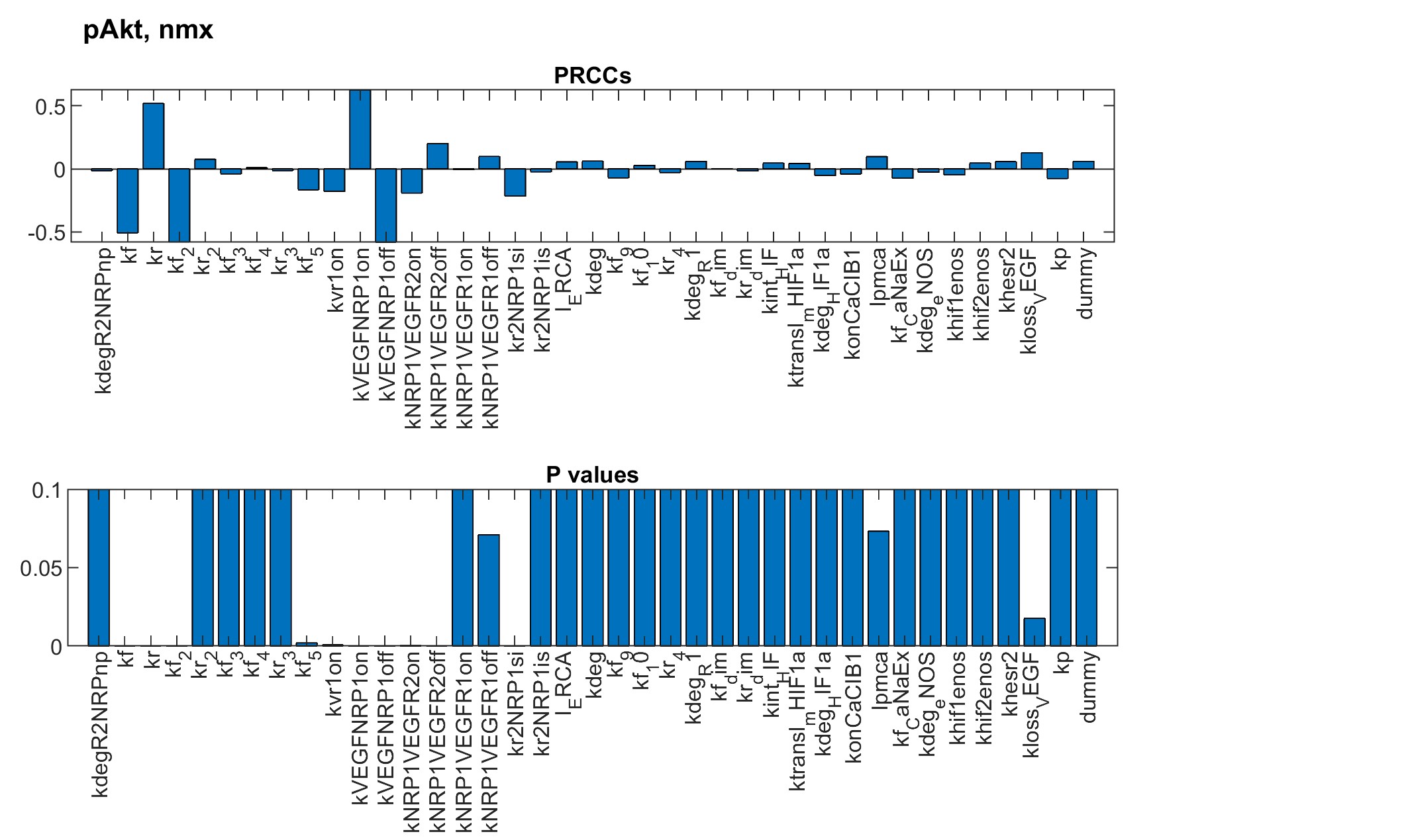

### Figure S11.jpg

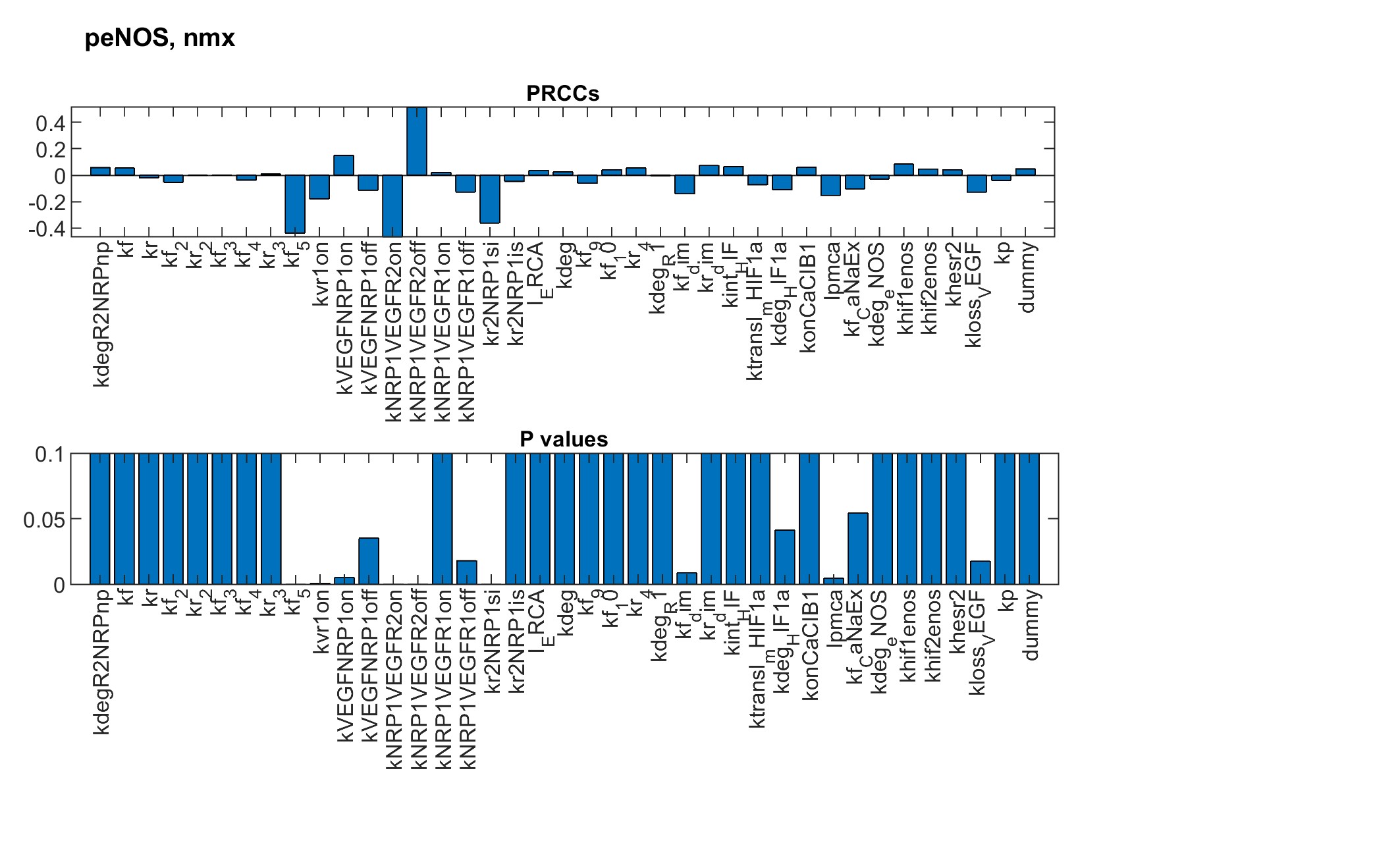

### Figure S12.jpg

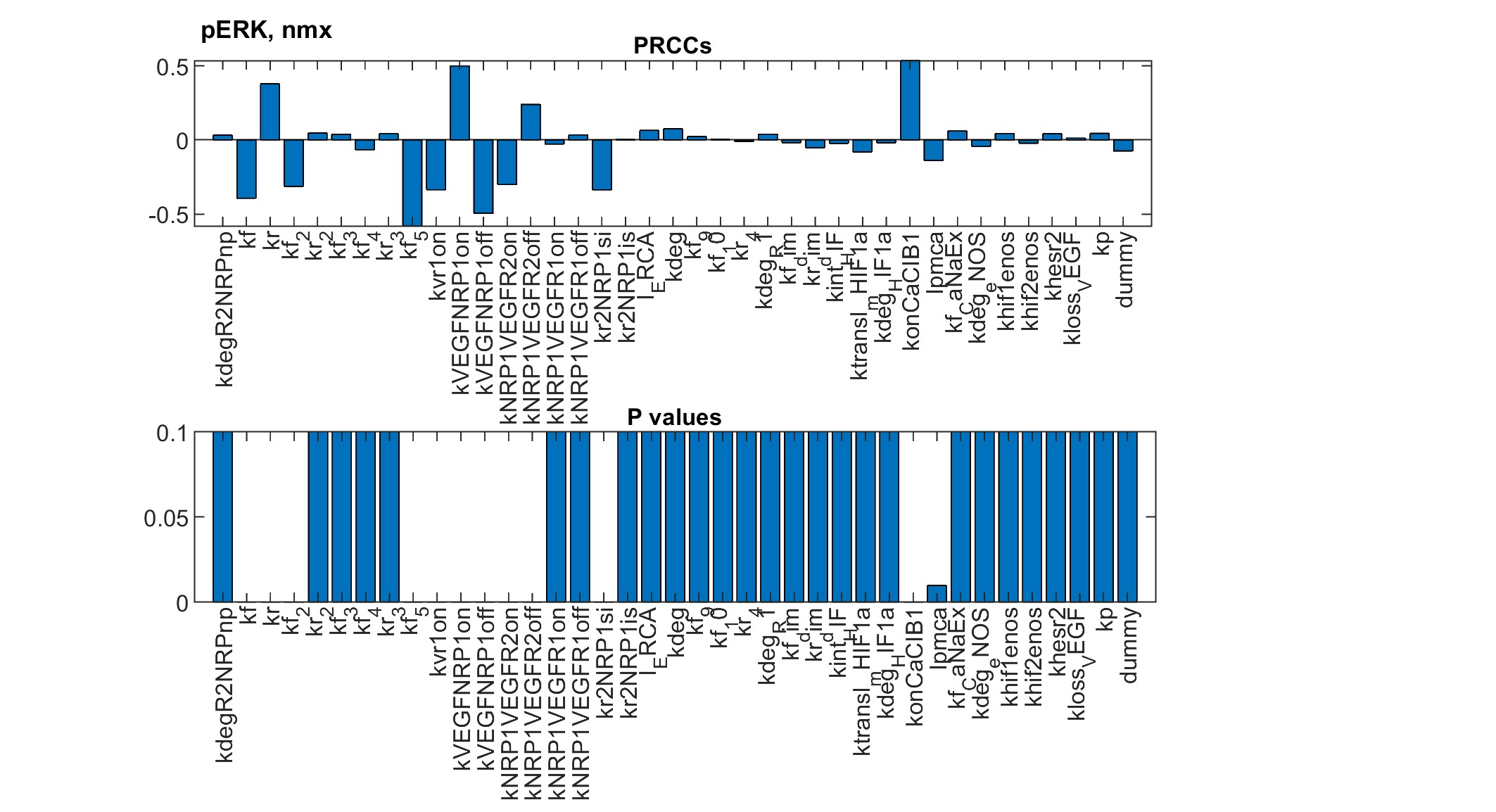

### Figure S13.jpg

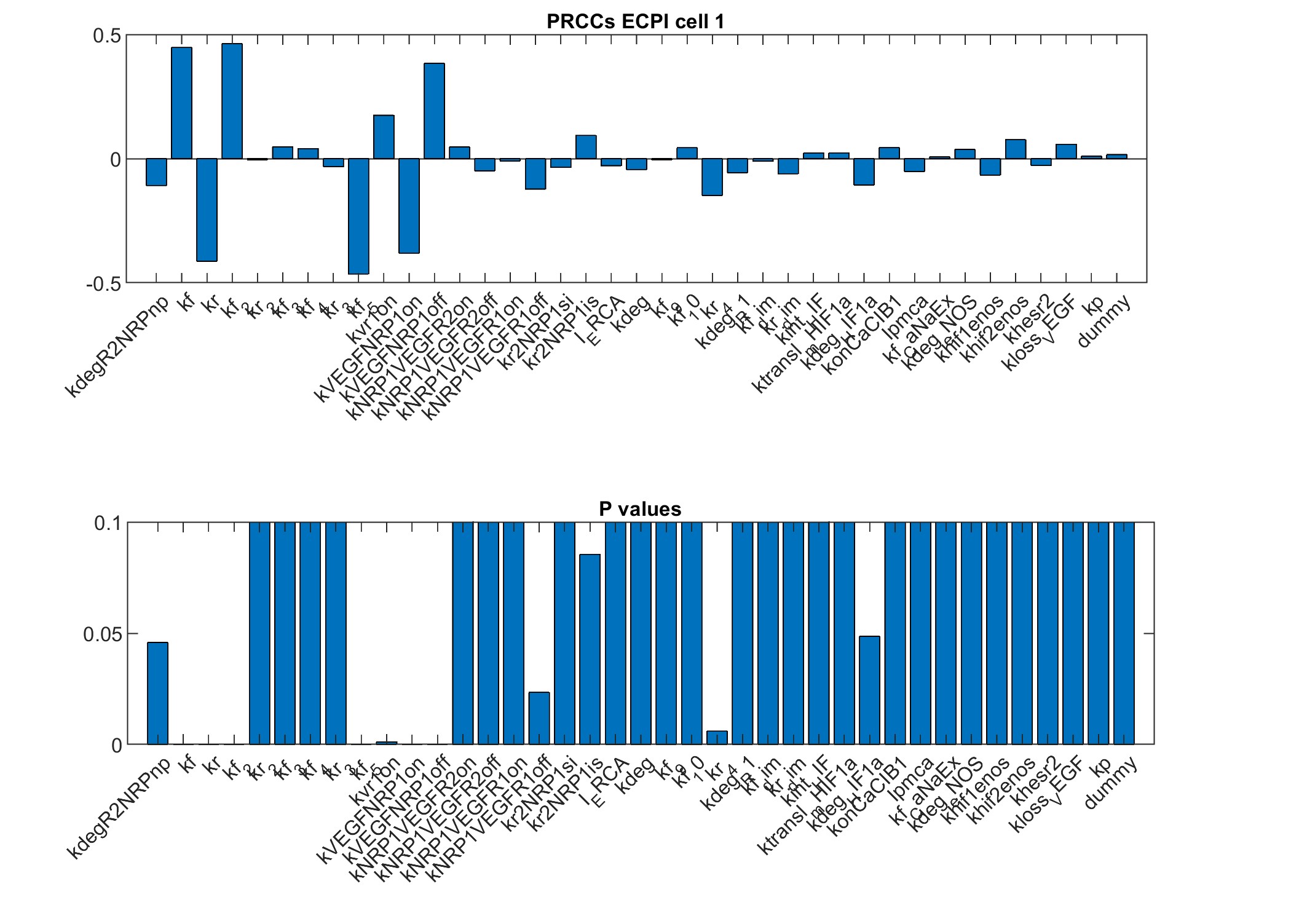

### Figure S14.jpg

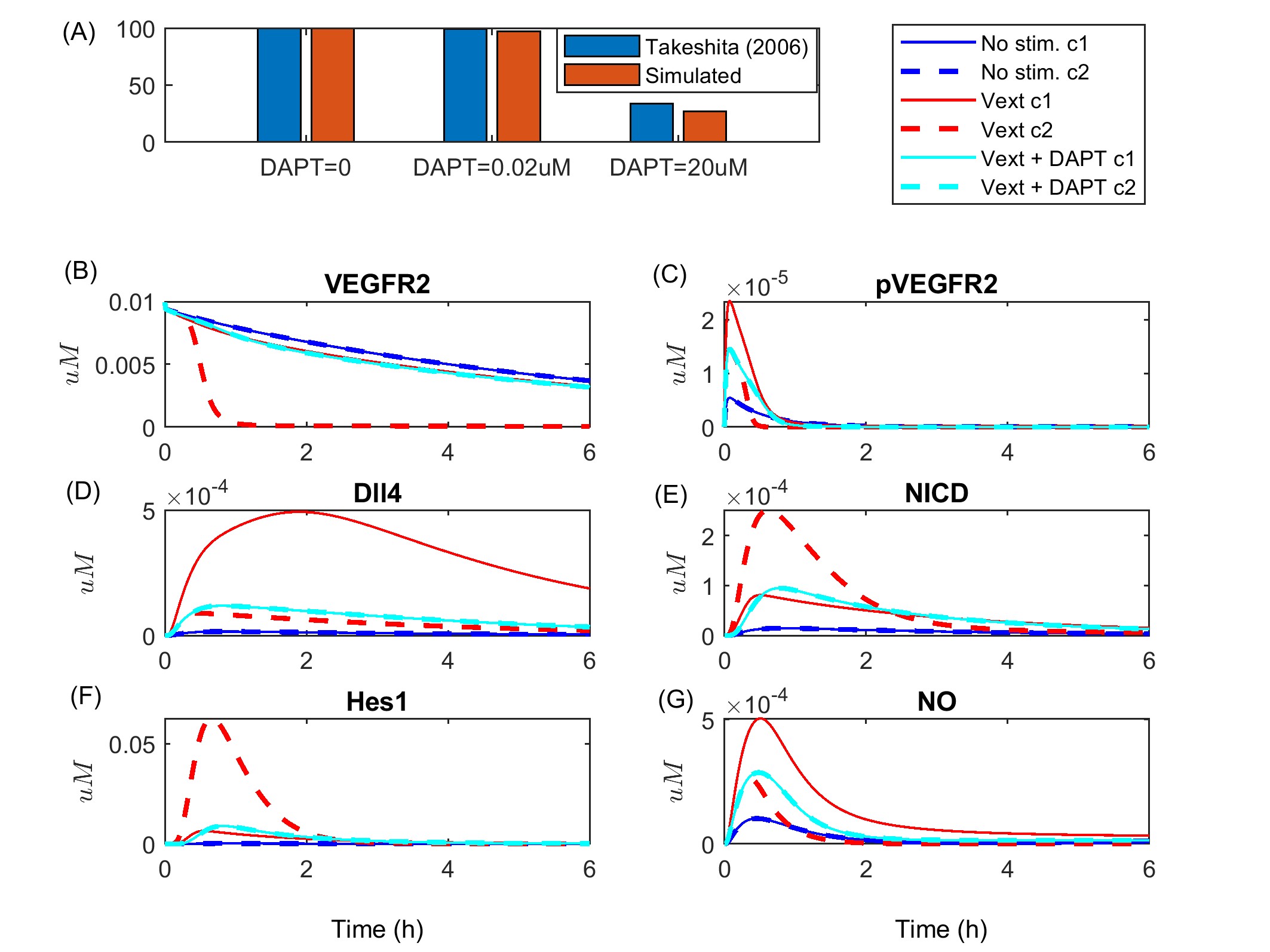
