## Supplementary Material for "Endothelial cells signaling and patterning under hypoxia: a mechanistic integrative computational model including the Notch-Dll4 pathway"

### 1 Signaling pathways description

In this section, we detail the signaling pathways included in the model and how they interact.

#### 1.1 The VEGF/VEGFR2 signaling pathway

In a hypoxic environment, VEGF is released to induce angiogenesis and normalize oxygen levels. Tip endothelial cells probe the environment with their filopodia protrusions and migrate to the site of higher VEGF concentration, where the ligands bind to VEGF receptors located on the cell's surface. VEGF can bind to VEGFR2, VEGFR1, or neuropilin-1 (NRP1), forming complexes with different reaction rates and binding affinities. The interaction with VEGFR2 leads to its phosphorylation. In the model, we also represent the internalization of free and bound VEGR2. The internalized receptors can be degraded or recycled to the surface. This signaling pathway is shown in Figure 2A. Additionally, we model the repression of VEGFR2 and stimulation of VEGFR1 by the Notch signaling pathway.

#### 1.2 The Akt signaling pathway

The Akt-eNOS signaling pathway is shown in Figure 2B. The pathway is initiated by VEGFR2 phosphorylation (surface and internalized). Phosphorylated VEGFR2 leads to the activation of Src in a complex with TSAD. Subsequently, Axl-1 is phosphorylated by phosphorylated TSAD-Src and then undergoes a second auto-phosphorylation. The double-phosphorylated Axl-1 is recruited to the membrane and it activates PI3K. Following this, PI3K leads to the phosphorylation of PIP2 to form PIP3 (a process that PTEN can reverse). PIP3 then recruits Akt and PDK1 to the cell membrane. Following, mTORC2 kinase acts on recruited Akt, leading to its phosphorylation on serine 473 residue. Then, the recruited PDK1 phosphorylates Akt on its threonine 308 residue. Akt activity is also upregulated under hypoxia, which we assume is mediated by the HIF1/2-VEGFA signaling(Martorell *et al.*, 2009).

#### 1.3 The Notch signaling pathway

During sprouting angiogenesis, the extracellular Notch ligands (Jagged 1, 2; Delta-like ligands 1, 3, and 4) initiate the signaling by interacting with Notch receptors (Notch 1-4) on the surface of ECs. Once the binding occurs, a series of proteolytic events occur, mediated by disintegrin metalloproteinase domain-containing protein 10 (ADAM10) and the $\gamma_{secretase}$ enzymes. $\gamma_{secretase}$ cleaves the surface of the Notch receptor intracellular domain (NICD), releasing it in the cell cytoplasm. NICD translocates to the nucleus, where it binds to the recombination of signal-binding protein for the immunoglobulin kappa J region (RBPJ). The complex acts as a transcriptional activator, promoting the transcription of Notch target genes such as transcriptional repressors Hairy Enhancer of Split (Hes) and Hes-related with YRPW motif (HEY1) genes. These genes regulate EC patterning and tip EC formation, migration, and proliferation, to promote functional angiogenesis (Swaminathan *et al.*, 2022).

The Notch signaling pathway is shown in Figure 2C. In our model, we represent VEGFR2 phosphorylation as an inducer of Dll4 through ERK phosphorylation (Fish *et al.*, 2017). Under hypoxia, the expression of Dll4 is also upregulated in different cell types, including EC (Patel *et al.*, 2005). As we found no data suggesting an alternative mechanism, we model this reaction as mediated by the increase in VEGFA under hypoxic conditions. Sequentially, it causes an increase in ERK phosphorylation (Minet *et al.*, 2000) and, finally, in Dll4 phosphorylation. Despite Dll4 upregulation under hypoxia being a well-established concept, we could not find time-course data demonstrating such an increase for HUVECs. Therefore, for model fitting, we considered the data available for RF/6A cell line (Dong *et al.*, 2011). The data provided show a similar increase in HIF1$\alpha$ and VEGFA as seen in HUVECs, which leads us to assume a similar behavior in time dynamics increase of Dll4. Dll4 of each cell binds to the Notch1 surface receptor of the other cell, initiating the Notch signaling. This interaction leads to cleavage of the NICD by $\gamma_{secretase}$ on the cell membrane. The cleaved domain translocates to the nucleus, binds to RBPJ (omitted in the model for simplification), and upregulates the expression of Hes1. Hes1 then downregulates the surface VEGFR2 expression (Jakobsson *et al.*, 2010). To represent inhibition of the Notch pathway, we also include in our model a commonly reported $\gamma_{secretase}$ inhibitor, N-[N-(3, 5-difluorophenacetyl)-l-alanyl]-s-phenylglycinet-butyl ester, or DAPT, which we model as a competitive inhibitor of the enzyme, with a rate of inhibition fitted manually to promote NICD inhibition proportional to the reported experimentally (Takeshita *et al.*, 2007).

We also represent VEGFR1 upregulation by Notch signaling, considering this as a consequence of Hes1 transcription. Under hypoxia, a similar behavior takes place, in which VEGFR2 is downregulated and VEGFR1 is upregulated (Ulyatt, Walker and Ponnambalam, 2011). This could indicate an upregulation of Notch signaling through Dll4 induction by hypoxia as a mediator of VEGF receptors modulation under hypoxia. With this consideration, we include reactions for receptor regulation directly through Notch1 signaling and not as an additional reaction directly linked to the HIF pathway.

### 1.4 The HIF signaling pathway

The oxygen-sensing pathway is represented in Figure 2D.

Under normoxia, oxygen molecules form complexes with Fe-DG-PHD2/PHD3, leading to HIF1/2-α hydroxylation and subsequent polyubiquitination by VHL (von Hippel-Lindau ubiquitin E3 ligase), which marks them for degradation. Finally, HIF undergoes proteasomal degradation (Hashimoto and Shibasaki, 2015; Zhao and Popel, 2015). However, under hypoxia, both PHD activities are reduced, and the hydroxylation of HIF1/2-α does not occur. Under hypoxia, HIF1/2-α are stabilized and move to the nucleus, where they dimerize with HIF1-β. The newly-formed complex then bonds to the hypoxia element site on DNA, leading to the expression of different genes as a response to hypoxia, along with VEGF, erythropoietin (EPO), and glycolytic enzymes(Zhang *et al.*, 2018; Korbecki *et al.*, 2021). HIF isoforms are also regulated by FIH, but we do not focus on this reaction in our model, as was done in previous work (Zhao and Popel, 2015).

We build a simplified model of the HIF pathway, implemented based on several other models from the literature(Zhao, Isenberg and Popel, 2017; Jaśkiewicz *et al.*, 2022; Ferrante, Preziosi and Scianna, 2023). In our model, oxygen ($O_2$) levels are considered constant (not consumed by the reactions). We include the two main forms of HIF described in hypoxia-induced angiogenesis, HIF1$\alpha$ and HIF2$\alpha$ (HIFs, in general) which undergo similar processes. We consider the effect of PHD2 and PHD3 as regulators of HIF signaling. PHD2 inhibits both HIFs and is upregulated by HIF1$\alpha$. The two HIFs upregulate the activity of PHD3. HIF2 is inhibited by both PHDs. PHDs inhibition of HIFs depends on $O_2$ concentration (for normoxia, more inhibition, for hypoxia, less) (Ferrante, Preziosi and Scianna, 2023). Additionally, basal rates of production and degradation of HIFs mRNA and protein are included in our model (Jaśkiewicz *et al.*, 2022). Finally, we model the binding to HIF1$\beta$ (under hypoxia) of both HIFs, and the downstream upregulation of VEGFA mRNA and protein due to hypoxia (Zhao, Isenberg and Popel, 2017). We consider the amount of HIF1$\beta$ as constant in the model. Our modeling strategy allows us to represent HIF response to different $O_2$ levels and follow the time-courses provided in experimental data.

To calculate the molar concentration of $O_2$ to be included in the model for simulations, we consider the oxygen solubility in water at 37C of ~1.3 μM/mmHg (Qutub and Popel, 2006; Zhao and Popel, 2015) and the oxygen partial pressure under normoxia for HUVECs similar to the environment used in *in vitro* experiments (21% $O_2 \approx 160$ mmHg $\approx 209$ μM) and under hypoxia (1% $O_2 \approx 8$ mmHg $\approx 10$ μM).

### 1.5 The Calcium cycling and eNOS-NO signaling pathway

IP3 activation by phosphorylated pVEGFR2 is one mechanism that controls the calcium cycling dynamics. Figure 2E represents this process, which relates to the phosphorylation of eNOS. Activated IP3 diffuses into the cell and binds to IP3-sensitive calcium release channels on the endoplasmic reticulum (ER). The binding leads to calcium release from the ER and a consequent decrease in calcium concentration in this compartment. This decrease causes calcium release-activated calcium channels (CRAC) to open, allowing extracellular calcium influx. The action of the surface membrane PM pump and ER SERCA channels balances calcium concentration in the cytosol.

Under hypoxia, we include in our model the activity of the sodium-calcium exchanger (Na+ − Ca2+) proposed by Berna et al. (Berna *et al.*, 2002). Hypoxia inhibits mitochondrial oxidative phosphorylation, which increases the cellular requirement of glucose for anaerobic glycolysis to replenish energy. The required glucose can enter the cell via a Na+-glucose-like co-transporter. The co-transporter increases the amount of Na+ in the cytosol, which is then balanced by the Na+ − Ca2+ exchanger, taking Ca2+ in and Na+ out. Through this mechanism, hypoxia increases intracellular calcium concentration. In the model, we present these mechanisms as a single reaction that depends on external Calcium concentration and HIFs concentration.

Calcium binds to Calmodulin and then to eNOS, leading to its phosphorylation (a process that also depends on Akt phosphorylation, activated by Src). Additionally, Src activates the chaperone protein HSP90, which facilitates eNOS phosphorylation. Phosphorylated eNOS reacts with its substrate Arginine, converting it to Citrulline and producing nitric oxide (NO) (Wu and Finley, 2020).

### 1.6 The Raf-Mek-ERK signaling pathway

The phosphorylation of VEGFR2 also activates the Raf-MEK-ERK pathway, as shown in Figure 2F. In this pathway, PLC-γ is activated by pVEGFR2 and leads to the generation of DAG and IP3. DAG binds to PKC and calcium, leading to a sequence of activations and phosphorylation of Raf, MEK, ERK, and sphingosine kinase 1 (SphK). Activated IP3 stimulates calcium release from the endoplasmic reticulum, and the released calcium can bind to calcium and integrin binding protein 1 (CIB), forming a complex. Active SphK (pSphK) is translocated to the plasma membrane by CIB, and a complex is formed between will Ca-CIB-pSphK. SphK1 will then phosphorylate its ligand (Sph), forming diffusible sphingosine 1 phosphate (S1P). Finally, S1P activates Ras forming RasGTP, which stimulates Raf phosphorylation. As previously mentioned, we model the increase in ERK1/2 under hypoxia as mediated by the increase in VEGFA transcription given HIF stabilization under hypoxia (Minet *et al.*, 2000).
